# Selecting Synthetic Data for Successful Simulation-Based Transfer Learning in Dynamical Biological Systems

**DOI:** 10.1101/2024.03.25.586390

**Authors:** Simon Witzke, Julian Zabbarov, Maximilian Kleissl, Pascal Iversen, Bernhard Y. Renard, Katharina Baum

## Abstract

**Background:** Accurate prediction of the temporal dynamics of biological systems is crucial for informing timely and effective interventions, e.g., in ecological or epidemiological contexts, or for treatment adjustments in therapy. While machine learning has proven its capabilities in generalizing the underlying non-linear dynamics of such systems, unlocking its predictive power is often restrained by the limited availability of large, curated datasets. To supplement real-world data, informing machine learning by transfer learning with synthetic data derived from simulations using ordinary differential equations has emerged as a promising solution. However, the success of this approach highly depends on the designed characteristics of the synthetic data.

**Results:** We suggest scrutinizing these characteristics, such as size, diversity, and noise, of ordinary differential equation-based synthetic time series datasets. Here, we demonstrate how to systematically evaluate the influence of such design choices on transfer learning performance. We conduct a proof-of-concept study on three simple, but widely used systems and four real-world datasets. We find a strong interdependency between synthetic dataset size and diversity effects. Good transfer learning settings heavily rely on real-world data characteristics as well as the data’s coherence with the dynamics of the model underlying the synthetic data. We achieve a performance improvement of up to 92% in mean absolute error for simulation-based transfer learning compared to non-informed deep learning.

**Conclusions:** Our work emphasizes the relevance of carefully selecting properties of synthetic data for leveraging the valuable domain knowledge contained in ordinary differential equation models for machine-learning based predictions. The code is available at https://github.com/DILiS-lab/opt-synthdata-4tl.

## 1 Introduction

Forecasting the development of biological systems over time and conditions enables, for example, the design of drugs and the implementation of effective measures against infectious diseases. However, this task is inherently difficult due to highly nonlinear dependencies present in biological systems, high measurement inaccuracies, and biological variability.

Besides traditional statistical approaches [1], machine learning (ML) has emerged to address non-linear forecasting settings [2]. However, ML requires large and curated datasets [3] to learn the dynamics of biological systems. The collection of data in these systems is commonly costly [4] or even unethical, for example, if responses to therapeutics are concerned. In other cases, such as early stages of infectious disease outbreaks, by definition, only few time points are available. Therefore, small biological data often pose a major challenge towards leveraging the capabilities of ML in biosciences [5].

Synthetic data have been used in biology to address the small data problem and for benchmarking purposes when ground-truth data is limited. For instance, Walonoski et al. [6] developed a software package for creating artificial electronic health records representing relevant statistical dependencies to support the development of downstream applications. We recently proposed the simulation of protein and peptide abundances considering peptide-protein graphs for benchmarking abundance imputation methods [7]. Carŕe et al. [8] simulated gene-regulatory networks, including the corresponding gene expressions. Furthermore, Matsunaga and Sugita [9] simulated molecular dynamics which they subsequently combined with experimental observations for prediction of conformational dynamics. More recently, Wang et al. [10] used a generative adversarial network (GAN) to produce synthetic protein features. Goncalves et al. [11] investigated the strengths and weaknesses of approaches such as probabilistic models and GANs for synthetic data generation, and they evaluated their findings using cancer data from SEER.

If information on system dynamics is available, mechanistic mathematical models can be used to create synthetic data. For example, Zhang et al. [12] used a mechanistic model to restrict the input space before applying ML to predict cell growth. A frequently used mechanistic model type are ordinary differential equations (ODEs). ODEs explicitly include and mathematically represent potentially intricate dependencies between variables and have been curated for various biological systems [13]. They are an established modeling method that captures the knowledge of domain-expert researchers who have intensively studied the corresponding dynamical biological systems.

ODEs have been combined with deep learning (DL) to estimate kinetic parameters for various tasks of system biology [14]. Costello and Martin [15] developed an ML model as a substitute for an ODE model to predict the derivative of metabolites over time from protein data. Further, ODE-derived synthetic data describing different facets of gene expression have been used for benchmarking [16–18]. Additionally, simulations from ODE models have been used to inform ML by including domain knowledge [19–22], for example, in biomedicine [23, 24].

One option for informing ML with ODE-based information is transfer learning (TL). TL was introduced as a means to transfer knowledge between related datasets to improve ML predictions [25]. In fine-tuning-based transfer learning, a (deep learning) ML model is pre-trained on data related to the task of interest (source domain data) to learn domain-relevant features [26]. The trained model weights then act as the initialization for fine-tuning the model on the target domain dataset, i.e., the type of data for which predictions should eventually be derived. TL has become a common technique for addressing small data issues and fully leveraging available datasets in biological domains. For instance, TL approaches improved infection [27–29] and drug response predictions [30].

Simulation-based TL, i.e., informing ML by pre-training with ODE-derived synthetic data, was used to address the issue of small data. Various applications have been proposed, e.g., in material sciences [31, 32], predicting lake temperatures [33], and improving predictions of chemotherapy treatments [34, 35]. As ODE solutions often describe the evolution of variables over time, they are particularly suited to deliver synthetic information for time series prediction tasks. Simulation-based TL helps leverage prior knowledge about biological systems by conditioning an ML model on exemplary trajectories of systems with similar dynamics before fine-tuning it to a specific task. This is useful not only in small data scenarios, but also when noisy data make it difficult for the ML model to learn the underlying relationships. To make simulation-based TL more broadly applicable and thus extend the application range of ODE models, we established and introduced the open-source Python library SimbaML [36] that simplifies the on-demand generation of diverse ODE-derived synthetic datasets along with ML-based training and inference.

However, the benefits of informing ML with ODE-based simulations depend on the qualitative and quantitative characteristics of the generated synthetic data [24]. For unsuitable settings, prediction performance can even worsen. Thus, consistently leveraging the predictive power of simulation-based TL for time series data is bound to a systematic investigation of such required characteristics of synthetic datasets. For this purpose, in this work, we design a pipeline that allows the multivariate investigation and selection of the characteristics of ODE-derived datasets with respect to their TL performance. Our approach demonstrates to a broader audience, such as the ML community, how the valuable systems biology knowledge contained in ODEs could be made available to ML-based time-series forecasting approaches. We highlight possible outcomes and usage of our pipeline by thorough investigation of three simple but widely researched biological systems, including the prediction of COVID-19 infections and predator-prey dynamics.

### Background for simulation-based TL and investigated synthetic dataset characteristics

Here, we consider simulation-based TL as a combination of ODE-derived synthetic data as prior knowledge for a deep learning model. This is particularly advantageous when target data are limited and critical dynamics have not yet been observed. In these cases, the amount of data is insufficient for pure DL prediction models to learn and forecast the novel behavior as their ability to approximate complex functions depends on rich and abundant data. Thus, prediction models pre-trained on theoretically possible system behavior, as in simulation-based TL, may outperform conventional DL. We deliberately focus on simple dynamical systems, DL models, and basic ODE calibration techniques to assess the applicability of simulation-based TL to the field.

We expect simulation-based TL to surpass pure ODE-calibration-based approaches when the underlying real-world process, and thus the observed data, diverge from the often simplified assumptions of the mechanistic model. Unlike ODE models, which are entirely governed by their mechanistic assumptions, TL approaches can retain flexibility and account for deviations from these assumptions while being informed by simulations [33]. This remains valid also when highly sophisticated parameter calibration and data assimilation strategies [37, 38] are applied to adapt the ODE model or system state to the data, such as ensemble Kalman filters [39]. We anticipate more refined data assimilation techniques to benefit both mechanistic models and simulation-based TL. This is because pre-training on synthetic data from ODE models that better match the observed data and the underlying real-world process will likely improve the pre-trained model’s predictions.

Thus, we expect simulation-based TL to excel in cases where there is not sufficient data to train a pure DL model, nor the mechanistic model is entirely correct. It is important to note that we position simulation-based TL as a complementary tool, not a replacement for existing time-series forecasting methods, with the potential to improve prediction performance in specific scenarios. We evaluate simple DL models commonly used in time series forecasting to represent their major architectural concepts [26]. This allows us to isolate and understand the influence of the pre-training performed during simulation-based TL. Our goal here is not to optimize absolute performance but to assess how specific design choices on the synthetic datasets affect simulation-based TL outcomes.

Like for traditional TL, the success of simulation-based TL depends on the appropriate selection of data from the source domain, as unsuitable source data can even degrade performance (“negative transfer”, [40]). Specifically, the simulated data should resemble the observed target data for simulation-based TL to be effective [31, 32, 34]. For example, Steinacker et al. [34] observed less accurate predictions if target data was showing sudden oscillatory dynamics or extreme behaviors that were not well-represented. We consider different degrees of coherence as an important factor in our selection of systems and datasets.

As further key characteristics of synthetic datasets used for pre-training, we focus on dataset diversity, size, and noise. The *diversity* of a synthetic dataset, that is, the variety of the dynamics represented within it, potentially influences the generalization ability of the DL model pre-trained on it. The broader the range of theoretically possible dynamics a DL model is (pre-)trained on, the less likely it is to overfit to specific (potentially incorrect) dynamics. At the same time, a more diverse set of training data could lead to less exact predictions for each specific setting and difficulties distinguishing between settings. Thus, both more and less diversity in the synthetic data could be beneficial.

Higher dataset diversity depends on the *synthetic dataset size* as more samples are necessary to introduce more variability. At the same time, for fixed pre-training settings, the synthetic dataset size also modulates how strongly the pre-trained DL models are conditioned on the synthetic data and deviate from their naive non-trained state. Thereby, larger datasets put more emphasis on the provided synthetic data and, in theory, enable the pre-trained model to better capture the dynamics of the synthetic data. As described above, this can be beneficial for simulation-based TL if the synthetic data fits the observed target data well, or detrimental to performance otherwise. Thus, we investigate different dataset sizes to discern these effects, and in order to enable representing increasing synthetic dataset diversity.

Noise plays an especially relevant role in biological time series, as these are commonly subject to pronounced fluctuations. We expect that adding *synthetic noise* to the synthetic data for pre-training improves simulation-based TL performance if noise appropriately mimicking the real-world processes is applied. At the same time, noise can dilute the temporal information and dynamics from the ODE model presented to the DL model for pre-training and thus might be detrimental to simulation-based TL prediction performance. Moreover, the addition of random noise to training datasets is a known strategy for enhancing the generalization capabilities of neural networks [41]. Thus, characterizing the effect of synthetic noise of different magnitudes is important. We consider both classical measurement noise and intrinsic noise. The latter is comparable to random effects modelled by stochastic differential equations [42].

Simulation-based TL, and especially our simulation approach of synthetic data, shares conceptual similarities with techniques like simulation-based inference, approximate Bayesian inference [38], and ensemble approaches of data assimilation [39]. These approaches rely on some form of prior distribution that is used to inform the subsequent steps. However, simulation-based TL only exploits these prior data to select the initial functional shape to start optimizing using the observed data. In Bayesian approaches, on the contrary, these guide the entire search for a posterior distribution. Further, in ensemble-based data assimilation, they serve for inferring new system states along with the observed data as updated initial conditions for the mechanistic model. Further, our aim is not to estimate the best-fit kinetic parameter of the ODE model (as is the goal in simulation-based inference) but to directly retrieve a forecasting function. In addition, we aim for an offline setup where training the forecasting function only relies on data up to a fixed point in time. As a result, a direct comparison between the methods would be inappropriate, and we therefore do not address it here. However, calibrated parameters derived from methods such as simulation-based inference could be used in the pure ODE-calibration-based approach as well as for improving the synthetic data in simulation-based TL.

## 2 Methods

### 2.1 Pipeline for Assessing ODE-derived Synthetic Data Characteristics for Transfer Learning

We design a pipeline to assess ODE-based synthetic dataset characteristics for simulation-based TL. Our pipeline consists of two main components. First, we generate synthetic datasets with different characteristics (Figure 1A-C): we consider the synthetic dataset size, the diversity of the ODE dynamics represented in the synthetic data, and the synthetic noise applied to the data. Second, we evaluate the benefit of each synthetic dataset configuration for a real-world target dataset in a simulation-based TL approach (Figure 1D-F): we pre-train deep learning (DL) models with synthetic data and employ the pre-trained models to train on and predict observed dynamics in the real-world dataset. In addition, we compare the simulation-based TL results to a baseline DL setting, in which DL models are exclusively trained on the real-world dataset without any pre-training. A second baseline is derived from calibrating the ODE system to the real-world dataset.

**Fig. 1.**
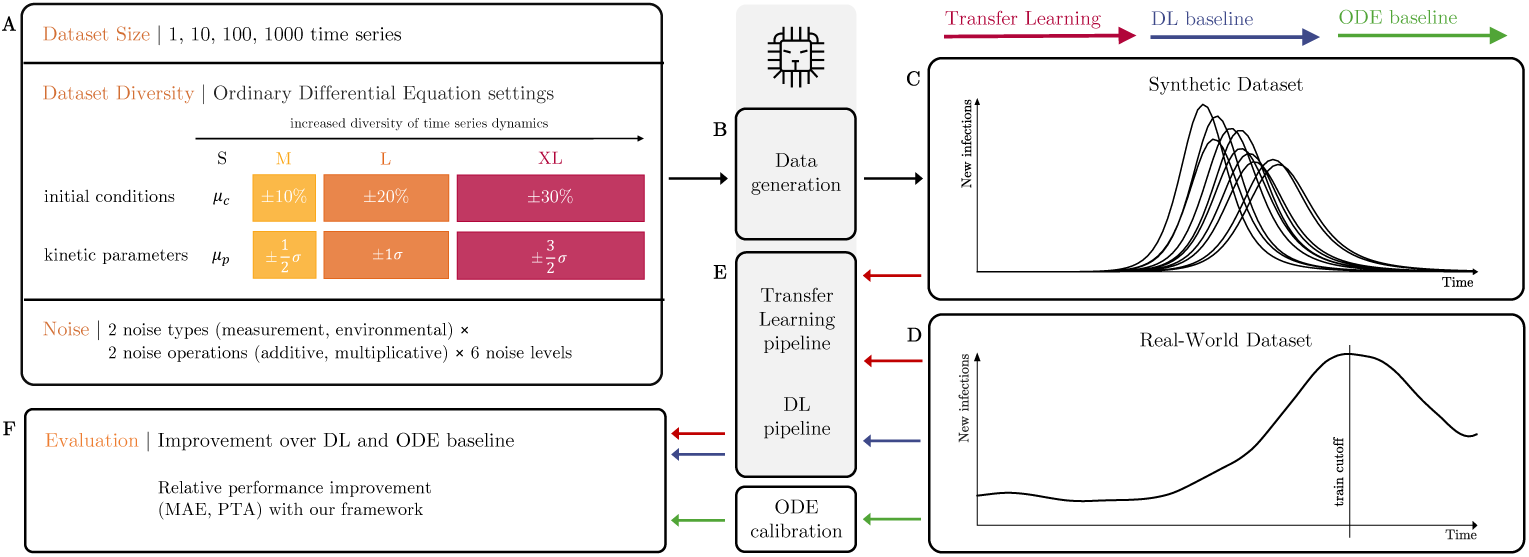
Overview of our pipeline. We propose scrutinizing the dataset size, diversity, and noise (A) to obtain suitable synthetic data for pre-training DL models. For an ODE model and a synthetic dataset configuration, we generate a dataset (B) by sampling uniformly from differently sized intervals for initial conditions and kinetic parameters, and by adding noise. Thereby, we obtain a dataset of one or more time series following the chosen ODE’s dynamic (C), which is then used for pre-training models before fine-tuning using (real-world) target data (D, E). We evaluate the performance of transfer learning with each specific dataset configuration against DL baseline and ODE calibration approaches conditioned only on real-world data (F). We use SimbaML [36] to generate synthetic time series and train DL models.

#### 2.1.1 ODE-based Synthetic Dataset Characteristics

To investigate the impact of synthetic dataset *size* on TL, we consider datasets with 1, 10, 100, and 1000 individual time series of fixed length. As pre-training epochs and other pre-training settings are fixed in all different configurations (see Supplement Section A.4 for details), this amounts to very slight pre-training on the dynamics of the chosen ODE model up to a strong conditioning on the ODE dynamics.

The *diversity* of a synthetic dataset influences the generalization ability of the pre-trained model. We propose altering it by two properties: (i) the variety in the initial conditions (ICs) of the ODE model, that is the starting values of the simulated timeseries, and (ii) the variety in the kinetic parameters (KPs) of the ODE model that enter the rate equations of the ODE system. To create a variety in these entities (and thus in the dynamics of the generated time series), their values are uniformly sampled from intervals during data generation. We translate a larger synthetic dataset diversity into larger ranges of the employed sampling intervals. Our pipeline considers four different intervals with increasing length (S, M, L and XL) for ICs and KPs around central values (see Supplement Section A.1.2 for details). The centers of the sampling intervals are derived from kinetic parameters and initial condition values that were estimated from related data sets in other studies. Overall, this approach amounts to a simple form of prior distributions. More complex prior distributions on these values are seamlessly compatible with our pipeline.

We further evaluate the impact of *synthetic noise* that is added to the ODE-derived data. In our pipeline, we support two different types of synthetic noise, either measurement or environmental, and two different noise operations, either additive or multiplicative. Measurement noise is the error that is expected to occur due to inaccuracies in the measurement process. We model measurement noise by adding or multiplying independent and identically distributed noise from a user-defined distribution (see Supplement Section A.1.3) to each variable at each time point of a synthetic time series. Environmental noise accounts for time-varying fluctuations in the environment, such as in enzyme activities, nutrient sources, and infectiousness of a virus, which directly impact the temporal development of the modeled entities. To capture the propagation of the effects of such fluctuations over time, we incorporate an environmental noise representation directly into the ODE model. We achieve this by adding a noise term to the rate equations, or by multiplying them by a noise factor at each time step. This is similar to the representation of randomness in sampled instances from stochastic differential equations with discretized time steps. For each of the described noise types and operations, we investigate different noise levels, where larger noise levels correspond to larger introduced perturbations (see Supplement Section A.1.3). The additive noise is thus scaled to fit the scale of the measured quantity. To reduce the combinatorial complexity, we investigate noise for the (fixed) combination of dataset size and diversity that yields the best simulation-based TL prediction performance. In addition, we always allow only one combination of synthetic noise type and operation at a time.

#### 2.1.2 Time Series Prediction, Baselines and Evaluation

We apply four commonly used DL model types in time series forecasting [26] in our simulation-based TL approach: long short-term memory neural network (LSTM), gated recurrent unit neural network (GRU), convolutional neural network (CNN), and dense neural network (DNN). Each model is combined with a two-layer perceptron that produces the final output.

For a specific synthetic dataset, we leverage SimbaML’s TL pipeline to pre-train each DL model on the ODE-based synthetic dataset. We then fine-tune the last two layers, i.e., the multi-layer perceptron, for five epochs on the train set of the real-world target dataset. Note that 10% of the synthetic and real-world time series in the training data are used as validation sets. To avoid overfitting during pre-training, all models are trained until the validation loss on the pre-training dataset ceases to improve. We use checkpointing throughout the complete pipeline to save the best DL model state according to validation loss. See Supplement Sections A.4 and A.6 for details on the DL model architectures and training setup such as dataset-specific train-test splits.

To evaluate the best prediction performance achieved with the different ODE-based synthetic dataset characteristics, TL results are compared to two baselines. First, we consider a non-informed, DL baseline setting in which we train the randomly initialized DL models only on the train set of the real-world data. Note that we compare the TL results of each DL model type (LSTM, GRU, CNN, DNN) to the performance of the corresponding architecture in the baseline DL setting. Second, we assess the performance of predictions via simulations of the ODE model after calibrating it, i.e., fitting its KPs, to the combined train and validation set of the real-world dataset (see Supplement Section A.7). Here, we intend to make the DL- and ODE-based approaches as comparable as possible by taking two measures. First, we match the overall complexities of the approaches by choosing straightforward ML-based models for time series forecasting (decisively more complex models like foundation models or transformer architectures could have been used) and equally straightforward kinetic parameter calibration methods (instead of more complex data assimilation strategies, as mentioned above). Second, we intend to compare approaches that employ real-world data for adapting (training or calibrating) the model only up to a certain point in time. This excludes complex DL-based (online) active learning strategies as well as some data assimilation strategies [39] for forecasting and the ODE calibration case whose performance may depend heavily on the characteristics of the dataset.

The time series predictions of the simulation-based TL, the DL baseline, and the ODE calibration are evaluated based on two metrics: mean absolute error (MAE) [43] and prediction trend accuracy (PTA), a variant of mean directional accuracy [44] (see Supplement Section A.5). While MAE considers the error between the predictions and the observed measurement in the real-world dataset, PTA only evaluates the correctness of the predicted trend, a desirable property for time series. We consider 1-PTA such that lower values are better for both metrics.

### 2.2 Coherence Assessment

To assess the coherence of time series dynamics between the investigated real-world datasets and our synthetic datasets, we perform an analysis based on multivariate dynamic time warping (DTW) [45]. DTW has been previously used to measure the similarity between the source and target datasets and subsequently choose a suitable source dataset to improve performance and avoid negative transfer [40]. In our case, we are interested in how well our ODE simulations match our observed target data. We sample time series from a synthetic dataset and compute their DTW distance to the corresponding real-world time series. We compare the obtained distance distribution to a baseline distribution of DTW distances solely between synthetic time series from the synthetic dataset (see Supplement Section A.3). Decisive differences to the baseline distance distributions (in the direction of larger differences) can hint at a lack of coherence. However, the DTW distance only provides relative assessments of coherence for similar datasets. Thus, we only use this measure to distinguish coherence levels of different datasets compared to the same synthetic data originating from one of our models. Another method of assessing (in)coherence, denominated as partial incoherence in Table 1, stems from targeted observations of violated temporal relationships between modeled entities in the dataset in parts of the time series (see Supplement Section A.3).

**Table 1.**
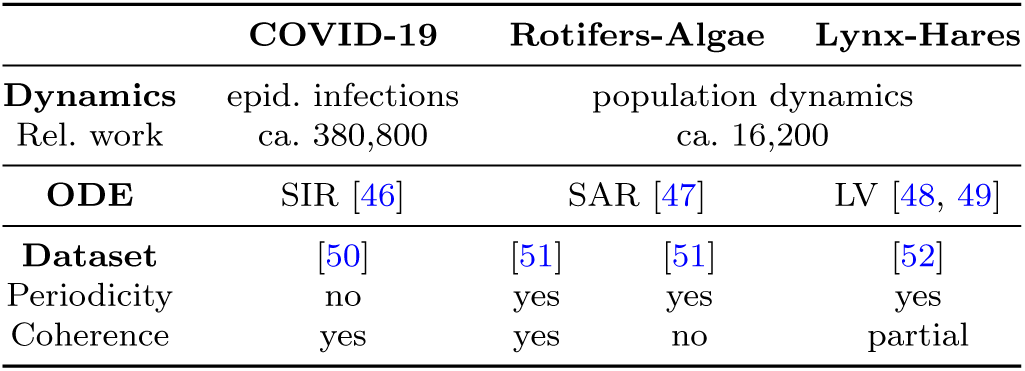
Overview of the biological datasets for which we scrutinize ODE-based synthetic datasets characteristics in a simulation-based TL approach. Related work (rel. work) captures citation counts of the biological dynamics from PubMed (see Supplement Section A.2.4). We indicate the ODE used to generate the synthetic datasets, the source of the real-world dataset, the periodic nature of the biological system, and if synthetic and real-world data in our experiments show coherent time series dynamics.

### 2.3 Biological Data and ODE Models

To demonstrate the output of our pipeline for assessing the impact of synthetic dataset characteristics for simulation-based TL, we select representative biological time series datasets using the following criteria: The set of datasets should (i) be publicly available; (ii) contain periodic and non-periodic time series among them as both are important types of biological dynamics; (iii) have been described to correspond to a specific ODE system; (iv) contain some dynamics coherent with the ODE and other dynamics being (at least) partially coherent and incoherent, ideally within the same biological system to minimize confounding factors.

Accordingly, we select four real-world datasets and forecasting problems from three biological systems of two distinct, widely researched biological fields (see Table 1 for their properties and Supplement Section A.2 for details). First, we aim to forecast the number of new infections from the fourth COVID-19 infection wave in Germany [50]. We use synthetic infection waves generated from the susceptible-infected-recovered (SIR) model [46] for pre-training. Second, we predict the predator counts in population dynamics of a predator-prey ecosystem containing rotifers, algae, and substrate for the algae [51]. Synthetic time series are generated from the nitrogen-algae-rotifers (SAR) model [47], an extension of the Lotka-Volterra (LV) predator-prey model. Third, we aim at forecasting predator counts using real-world predator-prey relations between lynx and snowshoe hares [53] with the original LV model [54] that we use for synthetic data generation.

Our coherence assessments (see Supplement Section A.3) confirm findings [51, 54] that the rotifers-algae dynamics in the investigated datasets contain predatorprey cycles incoherent with their respective ODE model dynamics (see Supplement Section A.2). Thus, we rely on two subsets of the observed rotifer-algae data as separate datasets (rotifer-algae coherent, rotifer-algae incoherent) to contrast dynamics that are coherent and (more) incoherent with synthetic time series. Further, based on previous observations [54] and hints from our DTW-based coherence assessments, we consider the lynx-hares dataset as partially incoherent. Since we do not have any indications for incoherence for the COVID-19 data and SIR-model, we consider it coherent.

For simplicity, we do not perform any calibration of the ODE but rely on literature-based KPs and ICs for the synthetic data generation with the ODE models. The kinetic parameter values, listed in Supplement Section A.2, were derived by fitting the ODE models to related datasets [47, 55, 56].

## 3 Results

We examine ODE-based synthetic dataset characteristics for simulation-based TL with our pipeline on four datasets. We first use our pipeline to evaluate the influence of synthetic dataset size and diversity on simulation-based TL by generating synthetic datasets with different combinations of numbers of time series and sizes of sampling intervals for KPs and ICs (Section 3.1 to 3.3), without any synthetic noise. We then address synthetic noise in a subsequent analysis (Section 3.4).

### 3.1 Beneficial Synthetic Dataset Characteristics

Our pipelines enables the identification of synthetic dataset and DL architecture configurations beneficial for simulation-based TL for time series forecasting. We compare its performance to the baselines of using DL without pre-training and calibrating the ODE model to the training data only. We show the predictions of each of these three approaches exemplarily for the lynx-hare dataset in Fig. 2. We find that our ODE-calibration-based predictions (based on the same training data as the DL-based models), as expected, cannot account for temporary switches in dynamics as caused by irregularities or other perturbations, exhibiting limited flexibility. Simulation-based TL with suitable synthetic dataset characteristics enables leveraging the valuable knowledge contained in ODEs and outperforms both baselines in all four datasets in terms of 1-PTA (Table 2). In terms of MAE, it outperforms the DL baseline for three out of four datasets and the ODE calibration-based results for two of the datasets (see Table 2 and Supplement Section A.8).

**Fig. 2.**
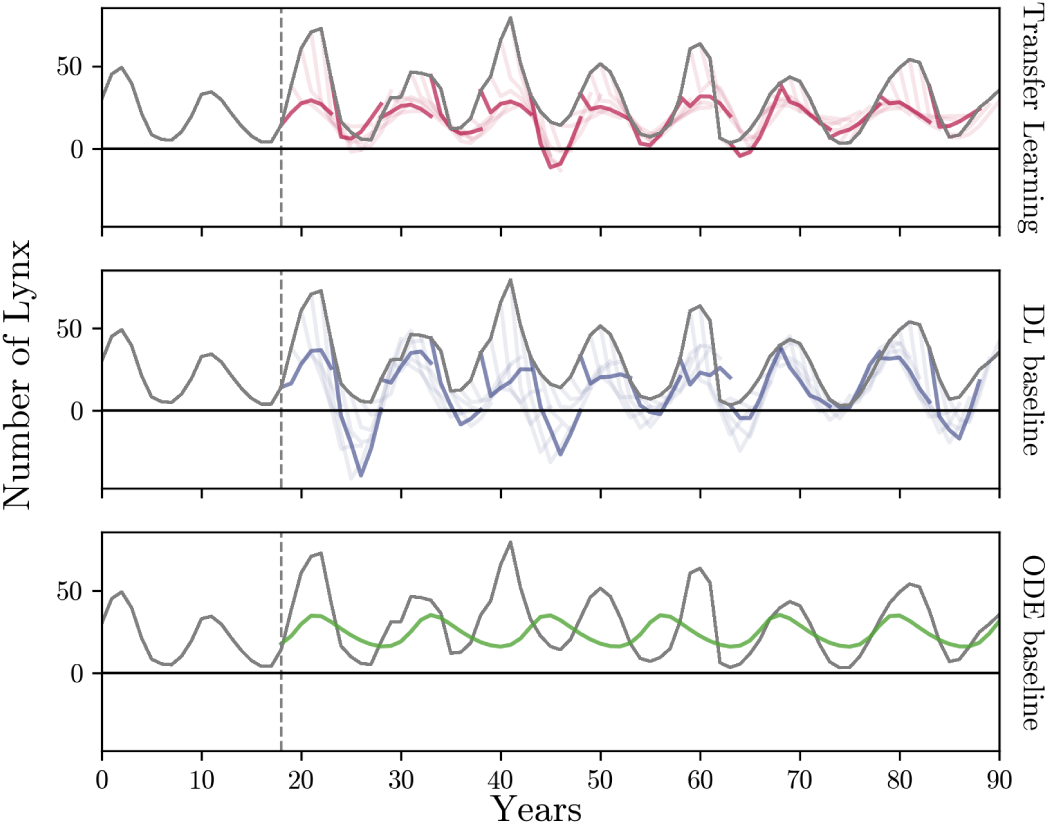
Time series forecasts for the lynx-hares dataset for the three prediction approaches. The predictions on the test set are shown for our simulation-based TL setting with beneficial synthetic dataset characteristics (red, top), the DL baseline (blue, middle), and the ODE baseline (green, bottom). The measured data are given in grey. As indicated by the respective vertical line, data from time steps 0-18 were used for training and validation or calibration (see Section 2). For both DL-based approaches, we show individual predicted time series for the number of lynx as colored lines. These predict five timesteps into the future, given the previous five timesteps of lynx and hare counts as input. As guidance, the lines start from the last input point. For better visibility, we only show every fifth prediction in full color. The predictions in the TL setting follow the ground-truth considerably more closely than the prediction of the DL baseline, where sharp drops from the last input occur regularly. Further, the TL setting produces much fewer insensible negative predictions than the DL baseline. The ODE baseline consists of a single time series after calibration. Due to the nature of the mechanistic model, predictions are always positive. However, irregularities in the dynamics cannot be accounted for (see time points 35-60 and [54]). Our simulation-based TL results reveal that combining ML and ODE modeling through synthetic data can counteract the shortcomings of each individual approach and yield superior results compared to employing both approaches separately. See Supplement Section A.9 for a visualization of the other datasets.

**Table 2.** Overview of the synthetic dataset characteristics most beneficial for simulation-based TL performance that we identified with our pipeline for each dataset (see Section 2.3). We report the number of time series (TS) and diversity of KPs and ICs for the best performing DL model types based on MAE and PTA (see Supplement Section A.8.1 for details on the selection criterion). The ‘DL’ and ‘ODE’ columns show the relative difference in MAE and 1-PTA of the most beneficial TL setting to the respective DL baseline run (with the same DL architecture as the TL), and the calibrated ODE. See Supplement Table S4 and S5 for more details on achieved performances.

| Dataset | Best TL Run |  |  | Model | MAE |  | 1-PTA |  |
| --- | --- | --- | --- | --- | --- | --- | --- | --- |
|  | TS | IC | KP |  | DL | ODE | DL | ODE |
| **COV-19** | 1000 | XL | XL | LSTM | -92% | -99% | -94% | -97% |
| **R-A (coh.)** | 100 | M | XL | LSTM | -23% | -3.8% | -44% | -47% |
| **R-A (inc.)** | 1 | S | S | LSTM | 5.6% | -60% | -25% | -8.3% |
| **L-H** | 1000 | M | S | CNN | -21% | -2.7% | -15% | -52% |

Thereby, superior performance of the simulation-based TL over a baseline DL approach depends on the coherence between time series dynamics in the real-world and generated datasets. With beneficial synthetic dataset configurations, simulation-based TL shows inferior average performance, by a very small margin (5.6% worse MAE), only in settings with pronounced incoherence between the real-world dataset and the synthetic data. In the case of the incoherent rotifers-algae dataset, our pipeline shows the best performance when pre-training on small synthetic datasets (a single time series). Thus, choosing the better-performing DL baseline setting would be advisable. For the COVID-19 dataset, we observe the greatest relative improvements with simulation-based TL over the DL baseline and ODE approach. Notably, any type of pre-training improves the prediction performance of the LSTM and GRU over the baselines for this dataset.

### 3.2 Impact of Synthetic Dataset Size

We can leverage our pipeline to systematically examine the influence of the dataset size on simulation-based TL performance (see Figure 3). For coherent time series dynamics between synthetic and real-world datasets (COVID-19, rotifers-algae coherent), the median MAE performance of TL runs can in general be improved by increasing the size of synthetic datasets. However, pre-training on 100 (and not 1000) time series yields the vast majority of best-performing dataset configurations for the (more) coherent rotifers-algae dataset. This might be due to subtle discrepancies between synthetic and real-world data. Also, the pre-trained DL model might struggle to generalize over too diverse time series from the larger synthetic dataset.

**Fig. 3.**
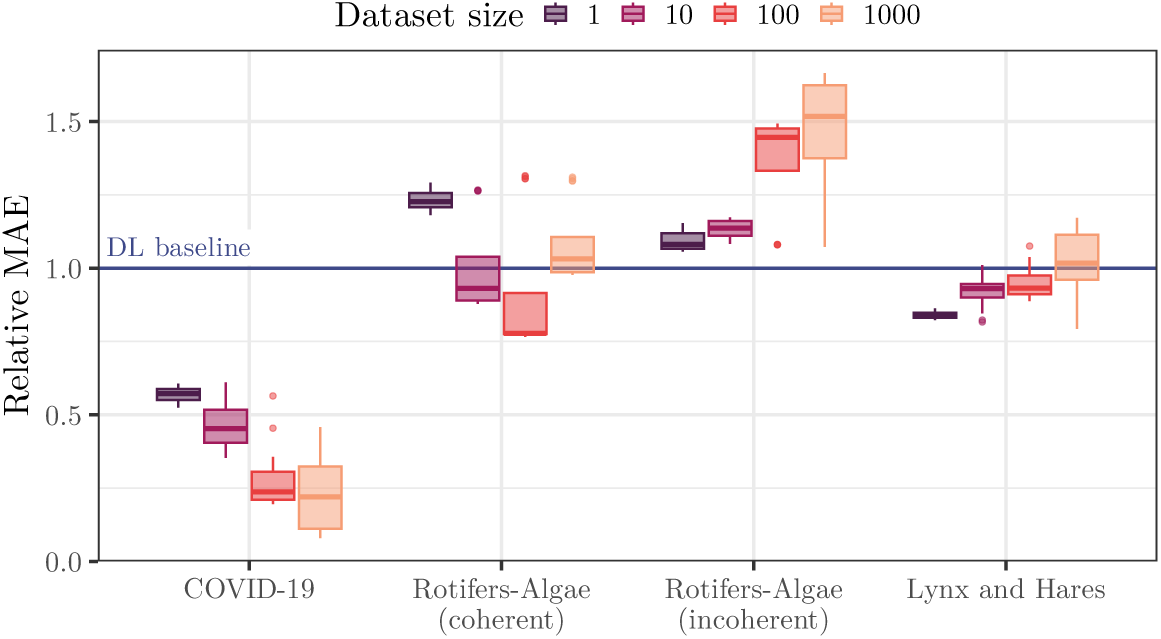
Univariate analysis of the performance impact of synthetic dataset size on simulation-based TL. The boxplots show the mean and five-fold standard deviations of the TL performance of various dataset diversity configurations (n=16) for each dataset size relative to the DL baseline (blue) for the best performing DL model architecture (see Table 2). Larger synthetic datasets improve MAE performance if time series dynamics in synthetic and real-world datasets are coherent (COVID-19, coherent rotifers-algae) and reduce it for incoherent datasets. We obtain similar results for PTA (see Supplement Section A.11, including ODE baselines that we do not show here due to their different scale).

For the incoherent rotifers-algae and lynx-hares datasets, our pipeline detects declining (median) performances for increasing dataset size and thus suggests TL synthetic dataset properties approaching the DL baseline setting. However, the variance of prediction performances for larger synthetic datasets across different diversity settings and DL architectures is substantially increased. Therefore, the largest dataset size setting (1000 time series) can yield both the best and worst performance results for a dataset, e.g., for the lynx and hares dataset, and careful characterization of the different configurations, as provided by our pipeline, is required.

### 3.3 Impact of Synthetic Dataset Diversity

The driver for the observed increase in variances in prediction performance with larger datasets is revealed in the multivariate analysis of dataset size and diversity (Fig. 4). We confirm that changing the dataset diversity by generating time series with a differing variety of KP has a particularly strong effect on prediction performance for larger dataset sizes. Univariate analysis shows that the absolute difference in MAE between the best and worst TL performance associated with each KP sampling interval correlates with its length (see Supplement Figure S14). MAE scores of TL are improved by increased diversity if the system dynamics between synthetic and realworld datasets are more coherent (COVID-19 and coherent rotifers-algae). For these datasets, we observe the most substantial improvement in MAE scores when increasing the dataset diversity by changing from the small (S) to medium (M) KP sampling interval. The opposite effect is observed in the incoherent rotifers-algae and lynx-hares datasets, where simulation-based TL performance decreases with increased dataset diversity, especially for larger synthetic dataset sizes. Due to the reduced coherence between synthetic and observed time series dynamics in these cases, the increased dataset diversity supports the conditioning of DL models on system behavior that is not contained in the target dataset.

**Fig. 4.**
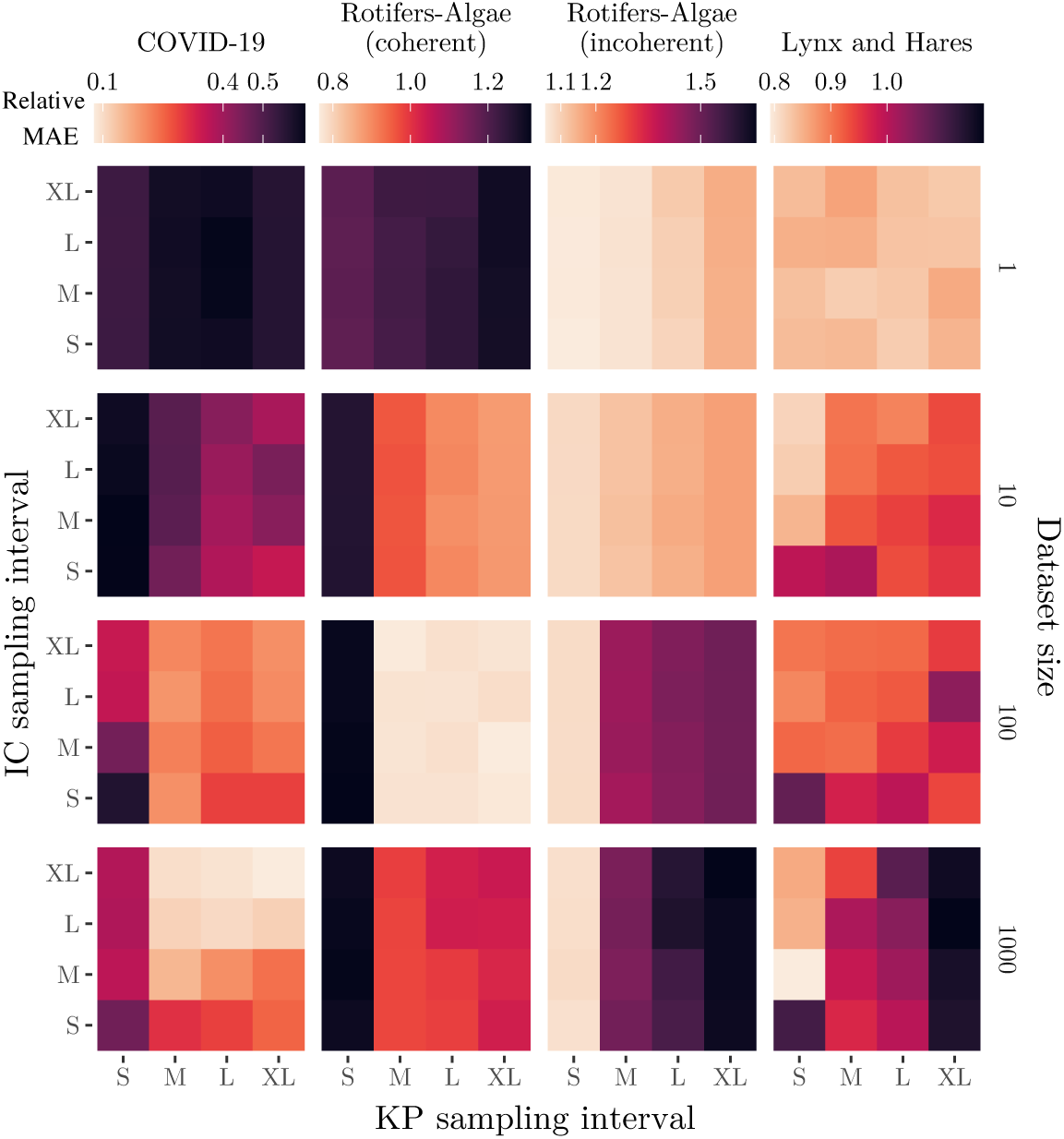
Multivariate analysis of the impact of dataset size, IC-based and KP-based dataset diversity on simulation-based TL performance. For each dataset, we show the MAE for the best-performing DL model architecture (see Table 2, ‘Model’ column) relative to its performance in the DL baseline. We observe that the most beneficial choice of the characteristics of synthetic datasets highly varies between real-world target datasets. This underlines the necessity of carefully characterizing ODE-derived synthetic datasets to fully leverage the predictive power of simulation-based TL. See Supplement Section A.10 for the figure with standard deviations and Section A.12 for the univariate analyses and PTA results.

Compared to KPs, manipulating dataset diversity by altering the IC sampling intervals shows decisively less influence on simulation-based TL prediction performance (see Supplement Figure S16). Different ICs often result still in similar time series dynamics, thus exerting a smaller influence on the overall diversity of the dataset. Nonetheless, in some cases, such as the COVID-19 and lynx-hares dataset, identifying a suitable IC sampling interval is still beneficial in a simulation-based TL setting (see Figure 4). For an investigation of how the synthetic dataset’s diversity impacts the PTA, see Supplement Section A.12.

### 3.4 Impact of Synthetic Noise in Synthetic Datasets

Further, we assess the impact of measurement and environmental synthetic noise on simulation-based TL performance for each real-world dataset. We use our pipeline to scrutinize the impact of the synthetic noise settings on simulation-based TL for the best performing setting (see Table 2). To make perturbations introduced by noise

comparable between datasets, we focus exemplarily on multiplicative noise here, i.e., noise is introduced as a factor drawn from a lognormal distribution around 1. We perform pre-training on synthetic datasets including either synthetic measurement noise or synthetic environmental noise. For each noise type, we consider six different noise levels corresponding to the degree of influence of noise on the data (see Supplement Section A.1.3 and Section A.14, also for additive noise results).

Using our pipeline, we find that small synthetic noise levels are beneficial for simulation-based TL for the datasets with coherent dynamics in synthetic and real-world target data (i.e., the COVID-19 and coherent rotifers-algae datasets, see Figure 5). In contrast, for the incoherent datasets, larger synthetic noise levels can increase performance up to baseline DL performance (for the rotifers-algae incoherent dataset) or can yield close to best TL performances (for the environmental noise and the lynx-hare dataset). For the lynx and hares dataset, increasing noise levels of multiplicative measurement noise reduces the prediction performance.

**Fig. 5.**
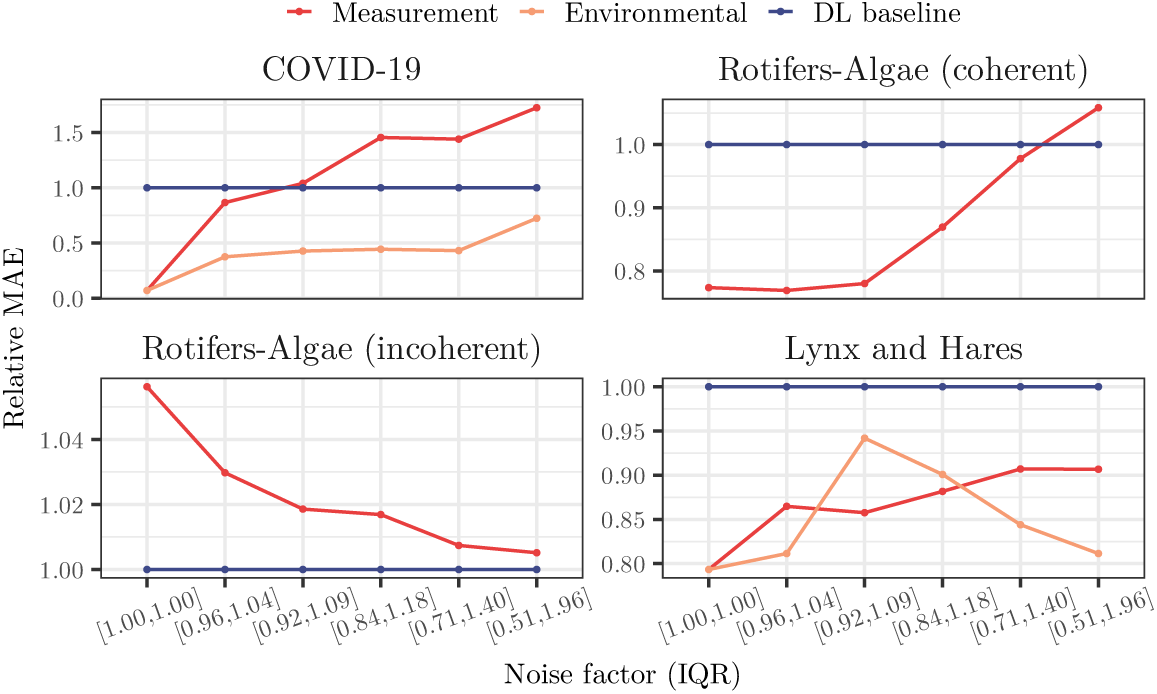
Univariate analysis of the impact of multiplicative synthetic noise on simulation-based TL performance. Lines show the mean MAE (over five seeds) of simulation-based TL performance relative to the DL baseline (blue) for each noise level indicated by the interquartile range (IQR) of the lognormal distribution used for sampling the noise factor. Environmental noise was not used for the rotifers-algae datasets due to numerical instabilities making simulations computationally infeasible. ODE baselines are omitted here due to their different scale, see Supplement Section A.14 for those and for the data variances. Low noise levels are beneficial for real-world datasets coherent with synthetic data, while larger noise levels can benefit simulation-based TL in incoherent settings.

The observation of low synthetic noise levels being beneficial for the COVID-19 dataset is in line with additional insights that we gained from synthetic-only experiments with our pipeline (see Supplement Section A.15). In these, we find that, in coherent settings, synthetic noise can especially improve performance when its levels are similar to those in the target dataset. Thus, the low noise levels in the COVID-19 dataset that arise due to the smoothing preprocessing steps (see Supplement Section A.2.1) are well matched with low synthetic noise levels achieving good simulation-based TL performance as observed with the help of our pipeline.

## 4 Discussion and Conclusion

We introduce a pipeline for investigating characteristics of ODE-generated datasets for simulation-based TL and apply it to a selection of widely researched biological systems and real-world time series datasets. We confirm the coherence in time series dynamics between synthetic and real-world datasets as an important factor that can determine the benefit of simulation-based TL over pure DL. Pre-training on larger or more diverse synthetic datasets, i.e., using broader IC and KP sampling intervals, together with low synthetic noise levels, can improve the performance of simulation-based TL for coherent settings. Deteriorating effects of high dataset diversity and large pre-training dataset size are observed when incoherence prevails, and increased synthetic noise levels can become beneficial. These findings are generally expected, but it is difficult to ascertain in absolute terms whether an ODE model is coherent or incoherent with the given dataset. Therefore, assessing the effects of synthetic dataset characteristics is required so that the knowledge from the ODE model can be optimally leveraged. Our pipeline provides assessment strategies, and the resulting simulation-based TL is regularly improved. If time series dynamics in synthetic and real-world datasets are incoherent and baseline performance is not reached, results from our pipeline naturally suggest the better suited non-TL setting.

The greatest relative performance improvement is achieved for the COVID-19 dataset. This is likely due to the small data setting where only parts of the non-periodic COVID-19 infection wave are available. As a consequence, baseline DL models are not able to infer non-observed dynamics and simple ODE calibration is observed to overfit to initial, non-representative system behavior here. With simulation-based TL, the DL model is pre-trained on expected but yet unobserved system behavior, e.g., a decline in infection numbers after the peak of the wave, which allows all TL runs to outperform the DL baseline, to a varying degree. This substantial advantage over the DL baseline setting is less present in our experiments with the periodic lynx-hares and rotifers-algae data. This might be due to complete oscillation cycles being already contained in the train set of the real-world data (see Supplement Section A.2). Therefore, we confirm that using simulation-based TL and our pipeline seems particularly beneficial in our considered small data scenarios when dynamic characteristics are incompletely observed. We do not consider more complex dynamics such as bifurcations or multi-stationarity [57] here. Furthermore, we neglect distribution shifts over time, such as those induced by genetic evolution in pathogens [58] or populations [59], or shifts in human behavior [60]. Whether our conclusions hold for processes with these more complex setups and richer dynamics remains subject to future investigations.

The phenomenon that increasing the diversity of datasets can improve DL model performance, and, in particular, its generalization abilities, has already been observed in other fields, such as computer vision [61]. Our pipeline enables detecting these cases and leveraging this fact also for simulation-based TL. Moreover, to fully leverage synthetic dataset diversity, the employed DL models in simulation-based TL have to be sufficiently large to capture (and reproduce) the distinct presented patterns during pre-training. Thus, the DL model sizes have to be taken carefully into account. Using our pipeline, we find consistent results when performing the same experiments using deeper and wider DL models (see Supplement Section A.11, Section A.12).

For the investigated dynamical systems, we observe that the diversity introduced by sampling ICs has a smaller influence on TL performance compared to those of the KPs. This reflects the overall lower influence of IC than KP on an ODE system’s dynamics here. However, ICs can have decisively more impact in more complex systems, e.g., containing conserved moieties or multiple steady states [57], and thus may still need to be considered carefully.

We also observe dependencies between dataset coherence and beneficial synthetic noise levels: If the synthetic time series captures the real-world characteristics well, large synthetic noise levels may obstruct learning informative patterns. In contrast, if synthetic dynamics are incoherent with the target dataset, increasing noise dilutes the (unsuitable) patterns in the provided dynamics, thus approaching an uninformed initialization with pre-training. Our pipelines enables detecting these cases and allows reverting to alternatives to simulation-based TL, such as the pure DL baseline, in incoherent settings.

### 4.1 Limitations

Our investigations shown here are limited to specified configurations for dataset size, diversity, and noise. Depending on the system of interest, they could be extended to examine broader or more fine-grained selections of values. Although we use simple prior distributions of kinetic parameters and initial conditions to sample synthetic data from in this work, our suggested characterization pipeline is compatible with more complex distributions such as obtained from calibration approaches. However, the effect of different choices regarding prior distributions remains the subject of future work. Particularly for the noise types, more complex settings could be more appropriate to represent real-world errors, e.g., combinations of measurement and environmental noise and different noise levels for different parts of the system. Furthermore, it could be of interest to study alternative ODEs for the datasets to investigate the effect of the complexity and suitability of the chosen ODE model. Similarly, other and more complex DL model architectures, such as GNNs [28, 62] may yield further benefits in a simulation-based TL setting. All these different variations could be readily investigated with the help of our presented pipeline.

Another limitation of our study is that we only take a simple approach to estimating the parameters of the ODE model for the ODE calibration baseline, and straightforward DL model architectures. The goal is to match the complexities of the approaches and to ensure comparability in terms of the same information for training the models. Both types of prediction models can be designed in a more complex fashion, which could boost their respective performances, e.g., by data assimilation approaches [38] or sophisticated transformer-based foundation models [63]. However, ODE-based and DL-based approaches inherently differ in the inference process where the DL-model uses input data from later time-points as well. Thus, overall, the comparison of our simulation-based TL to the ODE baseline in this proof-of-concept study is not optimal and should be interpreted with caution. Nevertheless, the observed decisive improvement of simulation-based TL over DL baselines stems from a well-controlled, highly comparable setting using identical DL architectures and input data, and thus remains valid.

It is of note that the degree of success of simulation-based TL highly depends on the availability of an ODE model that provides suitable dynamics for a real-world dataset. Although additional work is required to develop suitable ODE models or adapt published ones [13], we found that simulation-based TL with such models performed better than predictions based on non-informed DL. Our pipeline helps to detect cases where the ODE model does not suit the real-world target dataset that well (as in our incoherent settings), allowing data scientists to choose alternative prediction methods instead.

### 4.2 Outlook

Our pipeline and results presented could also help benchmark further datasets with ODE systems [64] and be extended to better leverage existing domain knowledge of more complex simulations, such as geographical spread of infections [65] or climate models [66]. Further, especially in these complex settings, it could be beneficial to compare or even combine simulation-based TL with other ODE-informed ML approaches. For example, customizing loss functions has been suggested to enforce modeled physical or biological constraints more strongly [33, 67]. This approach could also be combined with the estimation of kinetic parameters or even the functional form of the underlying ODE models within the DL model as has been suggested particularly for epidemiological use cases [68–70]. In addition, further alternatives to parameter-based TL could be worthwhile exploring [26].

## Supplementary information

Supplementary information is provided as a separate file.

## Declarations

### Ethics approval and consent to participate

Not applicable.

### Consent for publication

Not applicable.

### Availability of data and materials

The code is available at https://github.com/DILiS-lab/opt-synthdata-4tl. The COVID-19 dataset is available from [50], the rotifers and algae data stem from [51], the lynx and hares counts are available from [52].

## Authors’ contributions

JZ, SW, KB with the help of PI, MK, BYR conceived the study design. JZ performed the experiments and analyzed the data with the help of SW who also contributed the ODE calibration and coherence computation (together with PI) and MK (contributing the noise experiments). JZ, SW, PI, BYR, KB interpreted the data. SW, KB, BYR supervised the work. JZ drafted the manuscript with the help of MK, KB; JZ, SW, PI, BYR, KB substantially revised the draft. All authors agree to the final version.

## Acknowledgements

Not applicable.

