## Supplement for "Selecting Synthetic Data for Successful Simulation-Based Transfer Learning in Dynamical Biological Systems"

A.15 Synthetic-Only Experiments to Investigate the Relationship Between Noise in Synthetic and Real-World Data and its Impact on Simulation-Based TL Performance . . 28

### A.1 ODE-based Synthetic Dataset Generation for Simulation-Based Transfer Learning

We perform multivariate experiments that evaluate the impact of four different Ordinary Differential Equation (ODE)-based synthetic dataset characteristics on the prediction performance in a simulation-based Transfer Learning (TL) setting: the dataset size (see Section 3.2), the dataset diversity given by varying Kinetic Parameters (KPs) (see Section A.12.1, the dataset diversity given by varying Initial Conditions (ICs) (see Section A.12.2), and the synthetic noise supplied to the dataset.

#### A.1.1 Synthetic Dataset Size

To investigate the influence of the *size* of the synthetic dataset, we generate synthetic datasets by altering the number of time series in synthetic datasets. We define $Z = \{1, 10, 100, 1000\}$ as the set of synthetic dataset sizes. Each synthetic time series contains 100 equally spaced time points.

#### A.1.2 Synthetic Dataset Diversity Controlled by Initial Conditions or Kinetic Parameters

To alter the *diversity* of time series dynamics in the synthetic datasets, we define ICs and KPs sampling intervals of different lengths. We construct two sets of intervals $C$ and $P$ for ICs and KPs, respectively. Each set contains four elements that correspond to interval ranges of different lengths:

$$C = \{[\mu_c - i\sigma_{c,l}, \mu_c + i\sigma_{c,r}] \mid i \in \{0, 1, 2, 3\}\}, \tag{1}$$

$$P = \{[\mu_p - i\sigma_{p,l}, \mu_p + i\sigma_{p,r}] \mid i \in \{0, 1, 2, 3\}\}, \tag{2}$$

where $\mu_c$ and $\mu_p$ represent the interval midpoints. $\sigma_{c,l}$ and $\sigma_{c,r}$ define the step size with which the IC intervals are increased left and right from the interval midpoint $\mu_c$. The same applies for $\sigma_{p,l}$ and $\sigma_{p,r}$. The length of an interval is altered by the number of steps $i \in \{0, 1, 2, 3\}$ taken left and right from the interval midpoint. For an intuitive representation, we refer to each sampling interval according to its length, with S, M, L, and XL being smallest to largest.

To initialize the ODE models before data generation, the interval midpoints and step sizes are defined based on the literature, where the ODE models were fitted to a semantically similar dataset. Note that the step size taken left and right from the interval midpoint can differ depending on the type of calibration in each source. Details on the specific parameters of each ODE model used to generate the synthetic datasets can be found in Section A.2.

#### A.1.3 Synthetic Noise Types, Operations and Levels

Further, we investigate whether adding noise to synthetic data can enhance TL results. We consider two different types of synthetic noise (either measurement or environmental) and two different noise operations (either multiplicative or additive). Noise values are thereby drawn independently and randomly from a user-defined distribution $D$. The distributions that we use in our experiments are described below. In our experimental setup, they are normal distributions for additive noise and lognormal distributions for multiplicative noise to restrict noise factors to positive values, but other distributions are possible.

- We model *measurement noise* by adding or multiplying noise values element-wise to each time point for each variable of a synthetic time series.
- The *environmental noise* models time-varying fluctuations in the environment that propagate to subsequent time steps. They are implemented by changes in the ODE models' rate equations (derivatives). The noise values are constant for one timestep for solving the ODE system. We formally define the ODE system with additive environmental noise $\frac{d\hat{y}}{dt}$ as follows:

  $$\frac{d\hat{y}}{dt} = \frac{dy}{dt} + e_{\lfloor t \rfloor}; e_{\lfloor t \rfloor} \overset{\text{iid}}{\sim} D. \tag{3}$$

  The definition for multiplicative noise is equivalent, with the only difference being that noise terms are multiplied instead of added to the rate equations.

We investigate the influence of noise with six different *noise levels* on the data generation process. For multiplicative noise that we primarily consider here, we use a lognormal distribution with $\mu = 0$ and $\sigma \in \{0, 0.0625, 0.125, 0.25, 0.5, 1\}$. This results in the following interquartile ranges (i.e., middle 50%) for the randomly drawn noise factors that we use as noise level identifiers in the main text (Fig. 4): 1 (fixed values), $[0.96, 1.04]$, $[0.92, 1.09]$, $[0.84, 1.18]$, $[0.71, 1.40]$, $[0.51, 1.96]$. For the remainder of the Supplement, we refer to the noise level in increasing order according to $\sigma$.

### A.2 Real-World Datasets and Ordinary Differential Equations

In this section, we provide additional details on the ODE-based generation of synthetic datasets for a simulation-based TL approach to predicting the presented biological real-world datasets in Table 1 of the main text. We define IC and KP sampling intervals according to Eqs. 1 and 2, respectively. For all experiments, the KP sampling intervals are defined based on values that stem from fitting the ODE to semantically similar real-world datasets before simulating the synthetic datasets. Depending on the provided information by the source, the midpoints $\mu_p$ of the KPs' sampling intervals are set to the mean or median of the fitted distribution. If a confidence interval $[a_p, b_p]$ for each KP is available, the step sizes left and right ($\sigma_{p,l}$, $\sigma_{p,r}$) from the sampling interval midpoints are defined as follows:

$$\sigma_{p,l} = |\frac{\mu_p - a_p}{2}|, \qquad \sigma_{p,r} = |\frac{\mu_p - b_p}{2}|. \tag{4}$$

If the source provides no confidence intervals, the interval boundaries are increased by 10% increments from the mean or median:

$$\sigma_{p,l} = \sigma_{p,r} = 0.1\mu_p. \tag{5}$$

The same approach is used to define the step sizes with which the IC sampling intervals are increased:

$$\sigma_{c,l} = \sigma_{c,r} = 0.1\mu_c. \tag{6}$$

Details on the values used to define the interval midpoints $\mu_c$ of the IC sampling intervals in each experiment are provided in Sections A.2.1 - A.2.3.

#### A.2.1 COVID-19 and SIR Model

The real-world COVID-19 dataset used in the experiments is visualized in Figure S1. It contains new infections for each day from 2021-09-01 to 01-01-2022, i.e., 104 equally spaced time points. Note that the data is smoothed twice with a right-aligned window of size 7.

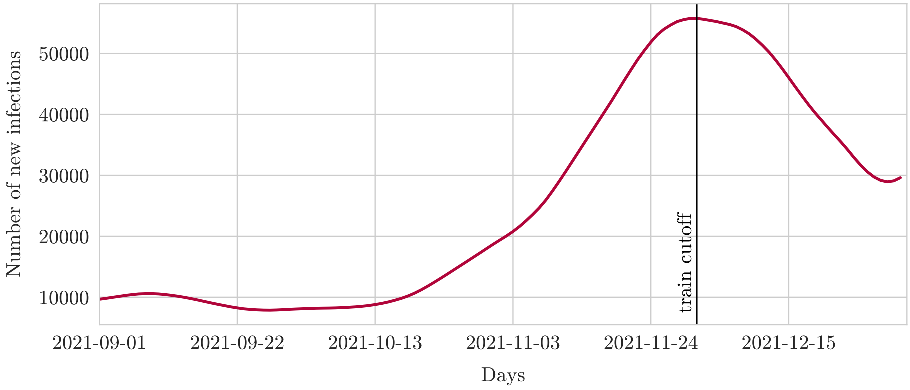

Figure S1: Infection dynamic of the fourth German COVID-19 wave (2021-09-01 to 2022-01-01) from a dataset by the Robert Koch Insitute [12]. The train cutoff is set to index 91, which is 2021-12-01. Consequently, index 90, i.e., the data from the 91th measurement, is the last predicted time point during training.

The synthetic datasets for the COVID-19 experiments are generated using the Susceptible-Infected-Recovered (SIR) model by Kermack and McKendrick [7]:

$$\begin{aligned} \frac{dS}{dt} &= -\frac{\beta SI}{N} \\ \frac{dI}{dt} &= \frac{\beta SI}{N} - \gamma I \\ \frac{dR}{dt} &= \gamma I, \end{aligned} \tag{7}$$

where $S, I, R$ are the modeled susceptible, infected, and recovered population, respectively. $N$ is the (constant) total population, $\gamma$ and $\beta$ are the kinetic parameters, and $t$ is the time. The ODE model's KPs and the number of initially infected people $I_0$ (used as midpoint $\mu_c$ in IC sampling interval for $I$) stem from fitting the model to a previous COVID-19 wave [4]. The parameters are listed in Table S1. The number of susceptible people $S$ is initialized with the population size of Germany, being $8.44 \times 10^7$ as of August 2023 [5]. This can be motivated by the fact that until the end of 2021, COVID-19 had long developed itself as a global pandemic, thus posing a risk to entire populations, including Germany. Due to emerging COVID-19 variants at the time, even previously infected people likely remained susceptible to an infection [14]. We use the development of daily new cases at time point $t_i$, $N_i$, for the synthetic datasets:

$$N_i = I(t_i) - I(t_{i-1}) + R(t_i) - R(t_{i-1}), \tag{8}$$

where $I_i$ and $R_i$ correspond to the number of infected and recovered people at time point $t_i$, respectively.

Table S1: Sampling interval midpoints $\mu_c$ and $\mu_p$, step sizes $\sigma_{c,l}$ and $\sigma_{c,r}$ for ICs $c \in \{S, I, R\}$, and confidence interval boundaries $a_p$ and $b_p$ of the fitted KPs $p \in \{\beta, \gamma\}$ [4] used to generate the synthetic datasets with the SIR model in the COVID-19 experiments.

| | $\mu_c$ | $\sigma_{c,l}$ | $\sigma_{c,r}$ |
|---|---|---|---|
| $S$ | $8.44 \times 10^7$ | $0.1 \times \mu_S$ | $0.1 \times \mu_S$ |
| $I$ | 19.2 | $0.1 \times \mu_I$ | $0.1 \times \mu_I$ |
| $R$ | 0 | - | - |
| | $\mu_p$ | $a_p$ | $b_p$ |
| $\beta$ | 0.41 | 0.32 | 0.51 |
| $\gamma$ | 0.12 | 0.08 | 0.18 |

#### A.2.2 Rotifers-Algae Data and SAR Model

The rotifers and algae data stems from a chemostats experiment by Blasius et al. [1]. The two investigated subsets in our experiments are shown in Figure S2. Subset A contains 33 measurements of rotifers and algae over 33 days. Their relationship follows the typical predator-prey cycle. This behavior is not visible in subset B, which contains 29 measurements over 29 days from the same experiment [1]. We predict the rotifers population in our experiments.

We use the Nitrogen-Algae-Rotifers (SAR) model as introduced by Rosenbaum et al. [13] to generate synthetic rotifer-algae dynamics. We generate 130 time points and use the last 100 time points to exclude the initial transient phase of the SAR model from the training dataset. The SAR model is defined as:

$$\begin{aligned}
\frac{dS}{dt} &= (S^* - S)\,\delta - \frac{1}{c_A}\frac{f_A S}{h_A + S}A \\
\frac{dA}{dt} &= \frac{f_A S}{h_A + S}A - \frac{1}{c_R}\frac{f_R A}{h_R + A}R - \delta A \\
\frac{dR}{dt} &= \frac{f_R A}{h_R + A}R - \delta R.
\end{aligned} \tag{9}$$

$\delta$ describes the inflow and outflow rate of the chemostat system, and $S^*$ refers to the concentration of nutrients in the inflow. $c_A$ describes the conversion of nutrients into biomass of algae. $c_R$ represents the transformation of algae into rotifer biomass. The growth rates of rotifers and algae are described by $f_R$ and $f_A$. $h_R$ and $h_A$ represent the half-saturation constants for rotifers and algae, respectively [13]. Rosenbaum et al. [13] fitted the ODE model's KP to data from a chemostats experiments different to our obtained real-world, target datasets from Blasius et al. [1]. The initial concentration of nitrogen $S_0$ (used as interval midpoint $\mu_c$ in the IC sampling interval for $S$) is set according on a simulated time series from the fitted ODE in [13]. The ICs' interval midpoints $\mu_c$ for algae $A$ and rotifers $R$ are defined based on their mean values in the chemostats experiment dataset

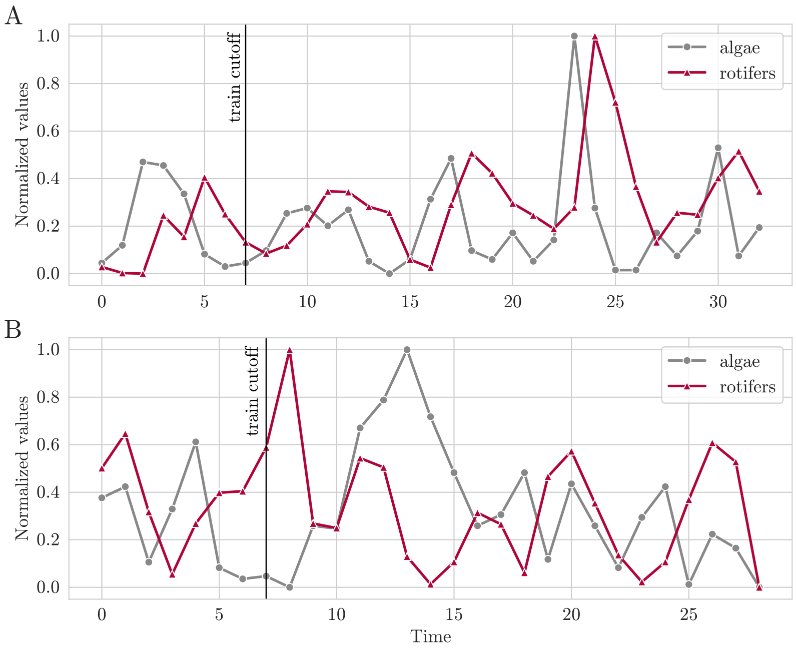

Figure S2: Real-world data from chemostats experiments with rotifers and algae [1]. We investigate subsets with coherent (A) and incoherent (B) system dynamics to those described by the SAR model (Equation (9)): Only in A do rotifer and algae populations show predator-prey cycles [1]. To simulate a data-scarce prediction scenario, the train cutoffs for both subsets are set to index 7. Consequently, index 6, i.e., the data from the seventh measurement, is the last predicted time point during training.

from which we define the two described subsets [1]. The parameters are listed in Table S2. Values for the other KPs of the SAR model, which are not listed in Table S2 are fixed as proposed by Rosenbaum et al. [13]: $\delta = 0.55\ \mathrm{d}^{-1}$, $S* = 80\ \mu\mathrm{mol\,L}^{-1}$, $h_A = 4.3\ \mu\mathrm{mol\,L}^{-1}$ and $h_R = 7.5$ cells/L.

Table S2: Sampling interval midpoints $\mu_c$ and $\mu_p$, step sizes $\sigma_{c,l}$ and $\sigma_{c,r}$ for ICs $c \in \{S, A, R\}$, and confidence interval boundaries $a_p$ and $b_p$ of the fitted KPs $p \in \{c_A, c_R, f_A, f_R\}$ [13] which are used to generate the synthetic datasets with the SAR model for the rotifers and algae experiments.

| | $\mu_c$ | $\sigma_{c,l}$ | $\sigma_{c,r}$ |
|---|---|---|---|
| $S$ [µmol L$^{-1}$] | 10 | $0.1 \times \mu_S$ | $0.1 \times \mu_S$ |
| $A$ [cells/L] | $0.52 \times 10^9$ | $0.1 \times \mu_A$ | $0.1 \times \mu_A$ |
| $R$ [ind/L] | $27.92 \times 10^3$ | $0.1 \times \mu_R$ | $0.1 \times \mu_R$ |
| | $ln(\mu_p)$ | $ln(a_p)$ | $ln(b_p)$ |
| $c_A$ [cells/µmol] | 17.68 | 17.60 | 18.76 |
| $c_R$ [ind/cells] | -11.82 | -11.71 | -11.93 |
| $f_A$ [d$^{-1}$] | 1.38 | 1.33 | 1.43 |
| $f_R$ [d$^{-1}$] | 0.23 | 0.20 | 0.26 |

#### A.2.3 Lynx-Hares Data and LV Model

The real-world, target datasets containing predator-prey dynamics of lynx and hares is visualized in Figure S3. It includes counts of yearly lynx and hare numbers from the MacKenzie River District, Canada, from 1845 to 1935, providing a time series of length 91. In our experiments, we predict the lynx population.

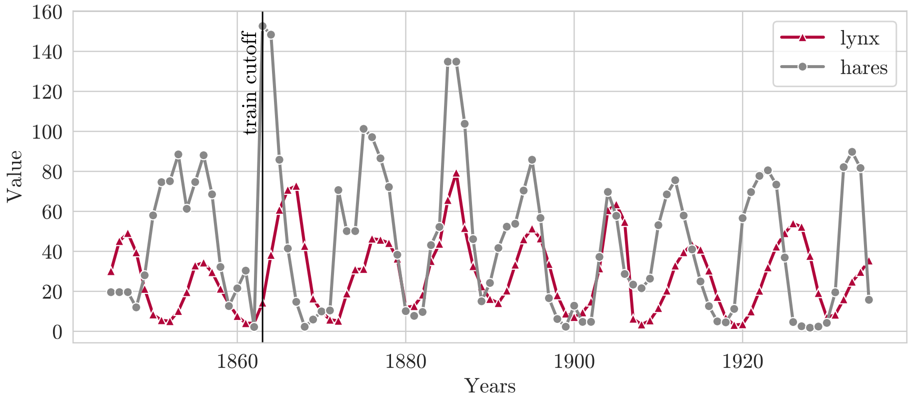

Figure S3: Populations of lynx and hares counted in the MacKenzie River District, Canada, from 1845 to 1935 [10]. The train cutoff (1864) marks the index of the train-test split that is investigated in the experiments. Consequently, the number of lynx from 1863, i.e. the 19th data point, is the last predicted time point during training.

We use the Lotka-Volterra (LV) model [9, 15] to generate synthetic data for the simulation-based TL approach. The LV model is defined as:

$$\begin{aligned} \frac{dH}{dt} &= \alpha H - \beta H L \\ \frac{dL}{dt} &= \delta H L - \gamma L. \end{aligned} \tag{10}$$

$L$ refers to the lynx and $H$ to the hare population. Values for IC and KP sampling intervals are given in Table S3. The listed parameters are the results of fitting the LV model to another lynx and hares dataset from Carpenter [3]. The sampling interval midpoints $\mu_c$ for $L$ and $H$ are set based on the mean of the lynx and hare populations in this dataset, respectively.

Table S3: Sampling interval midpoints ($\mu_c$ and $\mu_p$), step sizes ($\sigma_{c,l}$, $\sigma_{c,r}$) for ICs $c \in \{L, H\}$, and confidence interval boundaries $a_p$ and $b_p$ of the fitted KPs $p \in \{\alpha, \beta, \gamma, \delta\}$ [3] of the Lotka-Volterra model, which is used to generate the synthetic datasets for the lynx and hares experiment.

| | $\mu_c$ | $\sigma_{c,l}$ | $\sigma_{c,r}$ |
|---|---|---|---|
| $L$ | 20 | $0.1 \times \mu_L$ | $0.1 \times \mu_L$ |
| $H$ | 34 | $0.1 \times \mu_H$ | $0.1 \times \mu_H$ |
| | $\mu_p$ | $a_p$ | $b_p$ |
| $\alpha$ | 0.545 | 0.481 | 0.609 |
| $\beta$ | 0.028 | 0.024 | 0.032 |
| $\gamma$ | 0.803 | 0.711 | 0.895 |
| $\delta$ | 0.024 | 0.020 | 0.028 |

#### A.2.4 Relevance Assessment of Biological Systems

To quantify the relevance of the investigated biological datasets describing epidemic infection and population dynamics, we assess the numbers of results for suitable, advanced queries of the PubMed[1]-database:

- epidemic infection dynamics: The query '(epidemic infection predictions) OR (epidemiological time series) OR (SIR)' yields 380, 800 results.

[1] https://pubmed.ncbi.nlm.nih.gov/advanced/

- predator-prey population dynamics: The query '(predator prey) OR (Lotka Volterra)' yields 16, 245 results.

PubMed was queried on 2024-03-14.

### A.3 Evaluation of Coherence Between Synthetic and Real-World Data

We assess the coherence of the dynamics of synthetic and real-world data using a dynamic time-warping (DTW) distance. For multivariate time series, we use a single alignment for all species, which we establish with a multidimensional Euclidian distance. Therefore, incoherent lead-lag relationships result in high distances. The absolute value of this distance is not directly comparable between datasets. It depends on the length of the time series and the details of the dynamics.

We sample 1000 ODE-derived synthetic time series and calculate their DTW distance to the real-world time series to obtain a distribution of DTW distances. We then establish a baseline distribution by calculating the DTW distance between 1000 sampled pairs of synthetic time series. Figure S4 shows the histograms of the obtained distributions for each dataset.

We observe that one of the rotifers-algae dataset has an overall higher mean DTW distance of 7.40 than the other dataset, which has a mean of 7.03 and is, therefore, closer to the mean baseline DTW distance in the synthetic dataset (mean 5.12) (Figure S4A). Consequently, we term the dataset with smaller distances to the synthetic data coherent, while we term the other dataset incoherent (as a relative marker comparing it to the first dataset). The COVID-19 distances (Figure S4C) with mean 5.24 partially overlap with the baseline distances (mean distance 1.56). This hints at coherence between real-world and synthetic data. We do not observe any overlaps in the distance distributions for the lynx-hares dataset. Therein, the real-world dataset shows overall larger distances to its corresponding synthetic data than when comparing synthetic data only (mean distances 9.55, mean baseline distances 5.50) (Figure S4B). In addition with the previous observation of predator dynamics preceding prey dynamics in parts of the lynx-hares time series that contradicts classical predator-prey relationships [6], we consider the lynx-hare time series data as (partially) incoherent with the synthetic data of the Lotka-Volterra model.

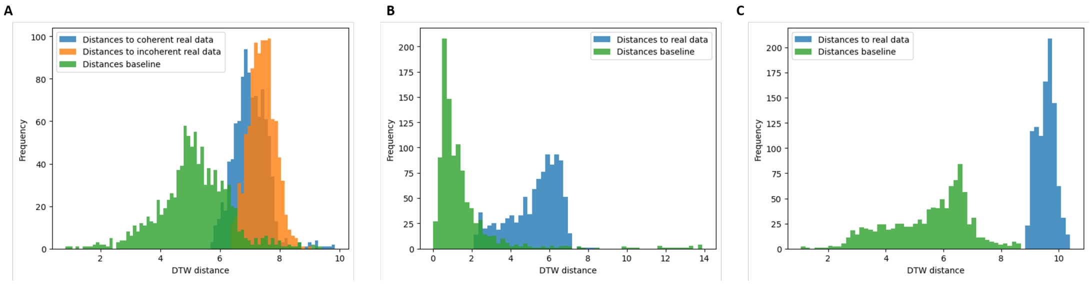

Figure S4: Distribution of dynamic time-warping distances between the sampled time series of the synthetic data set and the real-world data for (A) the coherent and incoherent rotifers-algae subsets, (B) the COVID-19 data, (C) the lynx-hares data. The baseline distributions are distances between random pairs of synthetic time series.

### A.4 Deep Learning Models for Simulation-Based Transfer Learning and Deep Learning Baseline

As described in Section 2.1.2, we employ four different Deep Learning (DL) model types in our experiments: Long Short-Term Memory Neural Network (LSTM), Gated Recurrent Unit Neural Network (GRU), Convolutional Neural Network (CNN), and Dense Neural Network (DNN). Figure S5 provides more details regarding their architecture and parameters. We consider two architectures of different sizes for each DL model type for our evaluation of the impact of synthetic dataset size and diversity. Investigations regarding the impact of synthetic noise on the TL performance with real-world data solely focus on the small DL architecture of the best-performing DL model type, as described in section A.6. Note that the DL baseline performance of each model type refers to predictions with the respective small DL architectures.

We apply univariate and multivariate time series data in our experiments. To handle both types of data, the `input_size` of the first LSTM [2] and GRU layer [3] of the LSTM and GRU DL models, respectively, is set to the number of input features. For the CNN, we set `in_channels` of the first `Conv1D`-layer[4] to the number of input features. For the DNN, we first flatten the input data before feeding them into the first fully connected layer. The size of the first DL layers is dynamically adjusted by considering the number of input features.

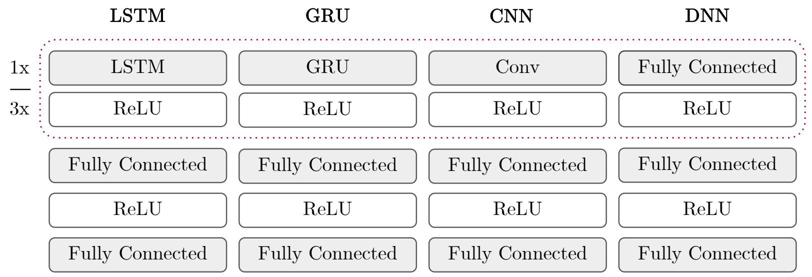

Figure S5: Overview of employed DL model architectures. The layer framed in red represents the models' feature extraction layer(s). The small architecture for each model type includes one feature extraction layer with 64 hidden states, filters, and neurons for the LSTM/GRU, CNN, and DNN, respectively. The large architecture comprises three feature extraction layers with twice the amount of parameters. The top model across model types and architectures contains 64 neurons per layer.

### A.5 Performance Evaluation Metrics

We evaluate the prediction performance using two complementary metrics. First, we consider the Mean Absolute Error (MAE) [16]. The MAE is a scale-dependent but easy-to-interpret error metric that is considered a standard evaluation method in time series forecasting. Lower MAE values represent a higher forecasting accuracy.

Besides the prediction error, we also aim to evaluate the forecasting direction. For this purpose, we introduce Prediction Trend Accuracy (PTA) as:

$$\text{PTA} = \frac{1}{n}\sum_{i=1}^{n} \mathbf{1}(\text{sign}(y_i - y_{i-1}) = \text{sign}(\hat{y}_i - \hat{y}_{i-1})), \tag{11}$$

where $\mathbf{1}$ denotes the indicator function and $n$ the length of the predicted time series. $y_i$ and $\hat{y}_i$ denote the ground truth and predicted value at time point $i$, respectively. In contrast to the more commonly used mean directional accuracy [2], the PTA does not penalize a shift between the predicted and ground truth values and instead fully focuses on the predicted trend. We achieve this by considering the predicted time point $\hat{y}_{i-1}$ instead of the ground truth $y_{i-1}$ as a reference when inferring the prediction direction of forecasts. For consistency across both metrics, we evaluate the results using 1-PTA. Therefore, smaller values indicate better results on all metrics.

### A.6 Experimental Setup for Synthetic Dataset Generation and Evaluation of Transfer Learning Predictions

For each of the four considered real-world datasets (see Section A.2 for more details), we perform a multivariate study to systematically evaluate the impact of synthetic dataset size and diversity

[2] https://pytorch.org/docs/stable/generated/torch.nn.LSTM.html
[3] https://pytorch.org/docs/stable/generated/torch.nn.GRU.html
[4] https://pytorch.org/docs/stable/generated/torch.nn.Conv1d.html

on the TL performance. For this purpose, we generate synthetic datasets without adding noise for each of the defined dataset size and diversity settings described in Section A.1. Thus, we, in total, consider $|Z \times C \times P| = 64$ synthetic dataset configurations. To ensure robust results, the synthetic data generation is repeated for five different seeds, which are also used during the subsequent TL experiments. Note that TL results for each synthetic dataset configuration were averaged over five seeds before evaluation and visualization.

For the best DL model type and optimal dataset characteristics (KP, IC sampling intervals, and dataset size, see Table 2 in the main text), we then investigate whether including noise in the synthetic datasets can further improve the performance. We systematically generate synthetic datasets using the six presented multiplicative levels (see Section A.1.3) for measurement and environmental noise and evaluate their impact on TL performance.

#### A.6.1 Time Series Prediction and Evaluation

For time series forecasting in the simulation-based TL and the DL baseline setting, we leverage the TL and DL pipelines in SimbaML [8].

For pre-training in the simulation-based TL setting, we provide the whole synthetic dataset as the train set. The synthetic time series are windowed using a rolling window with a shift of 1. The window size equals sum of input and output length for time series forecasts, which vary based on the dataset. The windowed time series are split into the first 90% for training and the last 10% for validation. Predictions on all windows of the train split contribute to the loss function. The DL models are trained on the train set until the loss on the validation set converges.

For fine-tuning in the simulation-based TL setting, we freeze the feature extraction layers of each DL model and solely train the fully connected layers for five epochs with the target datasets. We split the target dataset into a train set and a test set as described for each experiment in Section A.2. Again, time series in the train and test split are windowed using a rolling window with a shift of 1. We use 10% of the windowed train set as the validation set. All predictions on windows in the train set contribute to the loss function when fine-tuning the pre-trained DL model. Throughout the process, we use checkpointing to save the best model state based on validation loss for subsequent tasks.

Finally, we evaluate the trained DL model using MAE and 1-PTA (see Section A.5) on the test set of the target dataset. For that, we use a rolling window analysis[5] with a window shift of 1 to compute MAE and 1-PTA on the test set.

For the DL baseline, we only execute the steps as described for the fine-tuning part of the simulation-based TL setting. However, the number of training epochs and the used learning rate used for training DL models in the baseline setting can differ.

We define the following time series forecasting settings for our experiments (details on the mentioned experiment-specific time series forecasting settings are provided in sections A.6.2, A.6.3, and A.6.4):

- `epochs`: The number of epochs that the DL models are trained on the synthetic and real-world data for simulation-based TL and the DL baseline, respectively. The number of training epochs is defined differently in each experiment to ensure convergence of the training loss.
- `finetuning_epochs`: The number of fine-tuning epochs. These only apply to TL runs. In all experiments, we fine-tune DL models for 5 epochs using the train set of the real-world target data.
- `learning_rate`: The learning rate with which DL models are trained is defined differently in each experiment to ensure a smooth convergence of the training loss.
- `finetuning_learning_rate`: The fine-tuning learning rate is set to 1/10 of the `learning_rate` during pre-training. This only applies to the TL experiments.

[5] `https://de.mathworks.com/help/econ/rolling-window-estimation-of-state-space-models.html`, section 'Rolling Window Analysis for Predictive Performance' (accessed 2024-03-21)

- `batch_size` The batch size in all experiments is set to 64.
- `accelerator`: We set the accelerator to `auto` to automatically use a GPU if available. The TL experiments were performed using an NVIDIA A100 with 40GB of VRAM.
- `patience`: We set the early-stopping patience to 5 epochs during pre-training on the synthetic data during TL. The same applies to the DL baseline when trained on real-world data.
- `test_split`: The train-test split is defined differently in each experiment.
- `split_axis`: The split axis of the `test_split` to `vertical`. This way, we split each single time series according to the defined `test_split` (given by percentage) by their timestamps in two parts and use the first part with earlier time stamps as train set, the second part only for testing and evaluation. The splitting process is designed to avoid target leakage between the train and test set by ensuring that the last predicted timestep during training is not contained in the predicted time steps during testing.
- `input_features` and `output_features`: The input and output features are defined differently in each experiment.
- `input_length` and `output_length`: The input and output time steps vary between the experiments, while `input_length=output_length` holds true in all experiments.
- `loss function` As the loss function, we use PyTorch's `torch.nn.functional.mse_loss`-function, which computes the MSE of the windowed training data.

#### A.6.2 Experimental Setup for Simulation-Based TL and DL Baseline for Time Series Forecasting of COVID-19 Infections

During TL, we pre-train all DL models on the full synthetic datasets (split into train and validation sets), i.e., simulated time series of length 100, for 30 epochs using early stopping with a patience of 5. The learning rate is set to 0.001. Subsequently, the models are fine-tuned on the train set of the real-world infection numbers, i.e., the time steps 0-90 of the real-world time series (see Section A.2.1), for 5 epochs with a learning rate of 0.0001. In the DL baseline setting, DL models are trained for 20 epochs on the train set of the real-world infections with a learning rate of 0.001 and early-stopping using a patience of 5. For TL and DL, we use seven input time steps for the infected cases to predict the infection cases seven days into the future. For evaluation, we compute MAE and 1-PTA on the test set of the real-world time series, i.e., time points 91-104.

#### A.6.3 Experimental Setup for Simulation-Based TL and DL Baseline for Time Series Forecasting of Rotifers-Algae Population Numbers

The training parameters during TL are identical for both rotifers-algae datasets. We pre-train the DL model for 30 epochs on the ODE-generated data, i.e., simulated time series of length 100, with a learning rate of 0.001. The DL are fine-tuned on the train set of the real-world datasets, i.e., the time-steps with index 0-6 of each real-world time series datasets (see Section A.2.2), for five epochs, using a learning rate of 0.0001. For the DL baseline, all DL models are trained for 20 epochs on the same train set of the rotifers-algae populations as above, using a learning rate of 0.001 and a patience of 5. For TL and the DL baseline, we use three input time steps for rotifers and algae to predict the rotifer population three time steps into the future. For evaluation, we compute MAE and 1-PTA on the test set of the real-world time series, i.e., time points with indices 7-33 or 7-29 for the coherent or incoherent rotifers-algae dataset, respectively (see Section A.2.2).

#### A.6.4 Experimental Setup for Simulation-Based TL for Time Series Forecasting of Lynx-Hares Population Numbers

For TL on the lynx-hares data, DL models are pre-trained for 20 epochs on the synthetic datasets, i.e., time series with 100 time steps, with a learning rate of 0.01. The early-stopping patience is set to 5. We fine-tune the models on the train set of the real-world datasets, i.e., the time-steps with index 0-18, of the real-world time series (see Section A.2.3), for 5 epochs using a learning rate of 0.001. For the DL baseline, we train all DL models for 20 epochs using a learning rate of 0.01. In both prediction settings, we use five input time points for lynx and hares to predict the lynx population five days into the future. For evaluation, we compute MAE and 1-PTA on the test set of the real-world time series, i.e., time points with indices 19-90 (see Section A.2.3).

### A.7 ODE Calibration Baseline

For the ODE baselines, we use the full train set of the real-world datasets (i.e., including the 10% used for validation in DL, but not the separate test set) to calibrate each ODE system. This results in calibration sets of 91, 7, 7, and 19 time steps for COVID-19, coherent rotifers-algae, incoherent rotifers-algae, and lynx-hares, respectively. We rely on the *DiffEqParamEstim.jl*[6] and *DifferentialEquations.jl*[7] Julia packages for a simple calibration procedure. We use a Tsitouras 5/4 Runge-Kutta solver for the COVID-19 as well as Lynx and Hares datasets. Conversely, for the rotifers-algae datasets, we employ the library's capability to automatically switch from Tsitouras 5/4 Runge-Kutta to an Order 2/3 L-Stable Rosenbrock-W solver based on stiffness detection. For all experiments, we use the L2Loss provided by *DiffEqParamEstim.jl.* For the calibration optimization problem, we use the Broyden-Fletcher-Goldfarb-Shanno algorithm [11]. The result of this process is shown for each investigated dataset in Figure S6.

The real-world COVID-19 dataset [12] does not contain the susceptible, infected, and recovered populations, only the new cases. We, therefore, convert this time series to the series of cumulative cases and extend the ODE system by an additional compartment expressing the cumulative cases $C$ without influencing the development of the system:

$$\frac{dC}{dt} = \frac{\beta SI}{N} \tag{12}$$

We then obtain the number of new cases $D_i$ at timepoint $t_i$ by $D_i = C(t_i) - C(t_{i-1})$. For calibration, we use as ICs for variable $I(0)$ (and $C$) the first value of the train set (new cases at time point zero coincide with cumulative cases at this time point), a value of $8.44 \times 10^7$ for $S$ (full German population with negligent infections), for $I$ we use the first value from a dataset representing the new cases and a value of 0 for the variable $R$ (no recovery or removed at time point 0). Further, we restrict the parameter space of $\beta$ and $\gamma$ to $[0.00, 5.00]$ and $[0.00, 0.13]$, respectively.

Both rotifers-algae datasets only provide quantities for rotifers and algae but not for the nitrogen compartment of the SAR model. We, therefore, only fit those two compartments. As there are six orders of magnitude difference between the magnitudes of the values in the two time series (see Table S2 for fitted initial conditions for variables $A$ and $R$), we introduce weights to the loss function ($10^3$ for algae terms, $10^9$ for rotifer terms) that mitigate this difference to avoid prioritizing the fit to the algae time series. We use the first data point of the respective train set as ICs for rotifers and algae and default to the previously fitted initial value for the nitrogen species that has not been measured in the data (see Table S2). We set parameter bounds of $[0.00, 10^{30}]$ for all KPs.

For the lynx-hares dataset, we use the first values of the train set as ICs for the two modeled species and fit both populations. We restrict the parameter space for all KPs to $[0.00, 10.00]$.

For evaluation of the ODE calibration baseline, as predicted data, we use the ODE-derived time series obtained by solving each ODE with the ICs as used in the calibration procedure described above and the KPs obtained from the calibration procedure. Similar to the DL experiments, we window the resulting predicted time series with the dataset-specific window size and a shift of 1 before calculating the performance metrics by comparing it with the real-world target dataset.

We note that this procedure is to the slight disadvantage of the ODE baseline as the DL models are provided with more recent information as input. For the lynx-hares dataset, we perform an alternative evaluation approach where to obtain our predicted values, we always set the ICs to the

[6] `https://docs.sciml.ai/DiffEqParamEstim`

[7] `https://docs.sciml.ai/DiffEqDocs/`

last known value of a DL test sample and solve the ODE (with the calibrated KPs) for the five-time step horizon used during DL. Therefore, we obtain results that use similar information to the DL procedure when evaluating the ODE baseline. However, this approach is not straightforwardly applicable to the other datasets as we lack observations for at least one of the modeled species in the train and test sets. Thus, we restrict this evaluation to the lynx-hares dataset. In terms of performance on this dataset, we obtain similar but slightly better results for the MAE (14.72 compared to 15.29 with the first evaluation), but prediction performance is still comparable to the optimal simulation-based TL approach (MAE of $14.87 \pm 0.97$, see Table S4). For trend prediction as a performance assessment, the second evaluation is also slightly better than the first (1-PTA being 0.29 instead of 0.41 with the first evaluation), but optimal simulation-based TL outperforms both (1-PTA is $0.196 \pm 0.034$, see Table S5).

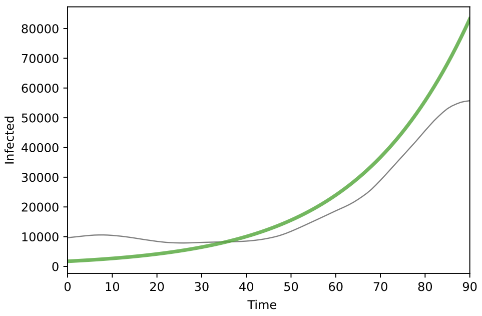

(a) Result of fitting the SIR model to the train set of the COVID-19 dataset.

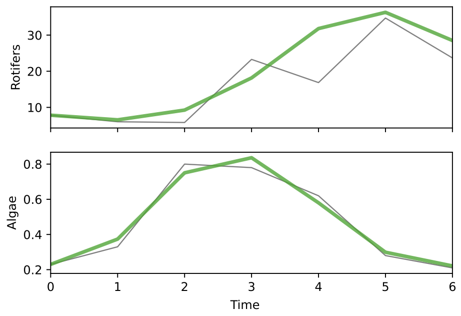

(b) Result of fitting the SAR model to the train set of the coherent rotifers-algae dataset.

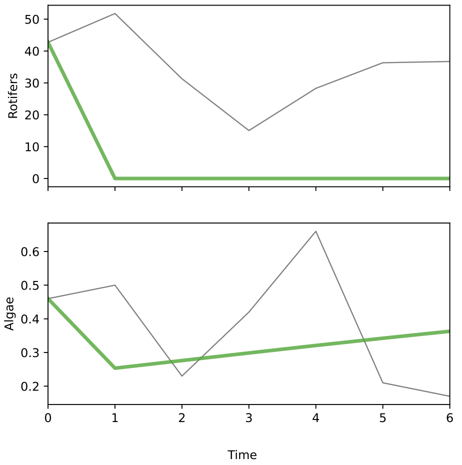

(c) Result of fitting the SAR model to the train set of the incoherent rotifers-algae dataset.

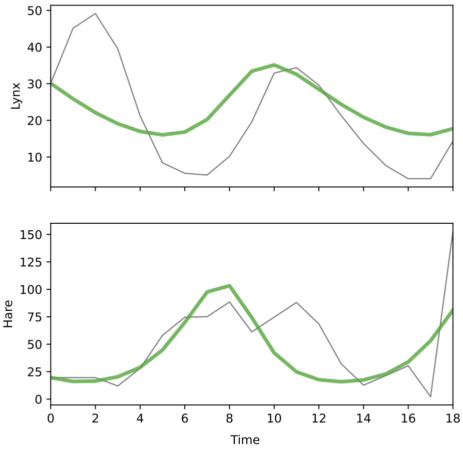

(d) Result of fitting the LV model to the train set of the lynx-hares dataset.

Figure S6: Overview of the resulting ODE baseline calibrations for each of the four investigated datasets.

### A.8 Optimal Synthetic Dataset Characteristics

#### A.8.1 Selection of Best-Performing Deep Learning Model Types

We determine the best-performing DL model type for each dataset according to the following optimization criteria: If one DL model type outperforms its performance in the DL baseline setting for MAE and 1-PTA for every synthetic dataset combination, this model is regarded as optimal. In the case that multiple models achieve this, the one with the best MAE performance is selected. If there is no DL model type that yields superior results than its performance in the DL baseline setting for all synthetic dataset combinations, the model type with the lowest MAE in the TL setting is selected. Note that for experiments on the two rotifers and algae real-world datasets, the same model is chosen to allow a better comparison of TL performance between the two datasets. Here, the optimal model selection process is done using the coherent rotifers-algae experiments.

Based on this optimization criterium and the raw data presented in Table S7 and Table S8 for MAE and 1-PTA, respectively, we identify the LSTM as the best-performing model type for the COVID-19 and both rotifers-algae experiments. On the lynx-hares dataset, the CNN provides the best prediction results. Details regarding the mean, median, standard deviation of the prediction performance for these models in the best TL setting are listed in Table S4. Of note, we find that the improvement of the simulation-based TL over the ODE baseline in the lynx-hare dataset (L-H) is smaller than the standard deviation of the performance estimates, suggesting that both approaches perform on par for this case.

Table S4: Overview of the optimal synthetic dataset characteristics for simulation-based TL that we identified with our framework for each dataset. We report the number of time series (TS) and diversity of KPs and ICs for the best performing DL model types based on MAE and PTA according to the presented optimization criterium in this section. We report mean, median and standard deviation (std) of the MAE over five runs with different initialization seeds for transfer learning ('TL MAE') and DL baseline ('DL MAE'), and the MAE for the ODE baseline ('ODE MAE' value). The 'rel.ch.' columns show the relative difference of the mean MAE of the best TL setting to the mean MAE of the respective DL baseline run (with the same DL architecture as the TL), and the MAE of the calibrated ODE.

| **Dataset** | **Coh.** | **Best TL Run** | | | **Model** | **TL MAE** | | | **DL MAE** | | | | **ODE MAE** | |
|---|---|---|---|---|---|---|---|---|---|---|---|---|---|---|
| | | TS | IC | KP | | mean | median | std | mean | rel.ch. | median | std | value | rel.ch. |
| **COV-19** | yes | 1000 | XL | XL | LSTM | 1532.89 | 1531.85 | 559.44 | 19379.78 | -92.1% | 19669.40 | 1400.16 | 1.08e5 | -98.6% |
| **Coh. R-A** | yes | 100 | M | XL | LSTM | 9.83 | 9.69 | 0.39 | 12.84 | -23.4% | 12.91 | 0.25 | 10.21 | -3.8% |
| **Inc. R-A** | no | 1 | S | S | LSTM | 13.03 | 13.01 | 0.12 | 12.34 | 5.6% | 12.32 | 0.12 | 32.22 | -59.6% |
| **L-H** | partial | 1000 | M | S | CNN | 14.87 | 14.83 | 0.97 | 18.75 | -20.7% | 18.77 | 0.85 | 15.29 | -2.7% |

Table S5: Overview of the optimal synthetic dataset characteristics for simulation-based TL that we identified with our framework for each dataset. We report the number of time series (TS) and diversity of KPs and ICs for the best performing DL model types based on MAE and PTA based on the optimization criterium in this section. We report mean, median and standard deviation (std) of the 1-PTA over five runs with different initialization seeds for transfer learning ('TL 1-PTA') and DL baseline ('DL 1-PTA'), and the MAE for the ODE baseline ('ODE 1-PTA' value). The 'rel.ch' columns show the relative difference of the mean 1-PTA of the best TL setting to the mean 1-PTA of the respective DL baseline run (with the same DL architecture as the TL), and the 1-PTA of the calibrated ODE.

| **Dataset** | **Coh.** | **Best TL Run** | | | **Model** | **TL 1-PTA** | | | **DL 1-PTA** | | | | **ODE 1-PTA** | |
|---|---|---|---|---|---|---|---|---|---|---|---|---|---|---|
| | | TS | IC | KP | | mean | median | std | mean | rel.ch. | median | std | value | rel.ch. |
| **COV-19** | yes | 1000 | XL | XL | LSTM | 0.033 | 0.026 | 0.021 | 0.538 | -93.8% | 0.506 | 0.110 | 0.981 | -96.6% |
| **Coh. R-A** | yes | 100 | M | XL | LSTM | 0.275 | 0.271 | 0.052 | 0.492 | -44.1% | 0.500 | 0.062 | 0.521 | -47.2% |
| **Inc. R-A** | no | 1 | S | S | LSTM | 0.415 | 0.425 | 0.014 | 0.555 | -25.2% | 0.575 | 0.041 | 0.452 | -8.3% |
| **L-H** | partial | 1000 | M | S | CNN | 0.196 | 0.184 | 0.034 | 0.231 | -15.0% | 0.232 | 0.008 | 0.408 | -51.9% |

#### A.8.2 Optimization of Transfer Learning Performance for PTA

Our developed framework allows the optimization of synthetic dataset characteristics with regard to MAE and 1-PTA. Table S6 lists the optimal selection of model type, dataset size, as well as ICs and KPs sampling intervals for obtaining the minimal 1-PTA. Note that this optimization does not follow the optimization criterium for the best-performing DL model type described in Section A.8.1.

Table S6: Overview of the optimal synthetic dataset characteristics for simulation-based TL that we identified with our framework when minimizing the 1-PTA for each biological dataset. We report the number of time series (TS), sampling interval length of KPs and ICs for the best performing DL model types regarding 1-PTA. We report mean, median and standard deviation (std) of the 1-PTA over five runs with different initialization seeds for transfer learning ('TL 1-PTA') and DL baseline ('DL 1-PTA'), and the 1-PTA for the ODE baseline ('ODE 1-PTA' value). The 'rel.ch.' columns show the relative difference in mean 1-PTA of the best TL setting to mean 1-PTA of the respective DL baseline run (with the same DL architecture as the TL), and the 1-PTA of the calibrated ODE.

| **Dataset** | **Coh.** | **Best TL Run** | | | **Model** | **TL 1-PTA** | | | **DL 1-PTA** | | | | **ODE 1-PTA** | |
|---|---|---|---|---|---|---|---|---|---|---|---|---|---|---|
| | | TS | IC | KP | | mean | median | std | mean | rel.ch. | median | std | value | rel.ch. |
| **COV-19** | yes | 10 | S | L | LSTM | 0.018 | 0.019 | 0.003 | 0.538 | -96.7% | 0.506 | 0.110 | 0.981 | -98.2% |
| **Coh. R-A** | yes | 10 | XL | L | GRU | 0.213 | 0.208 | 0.017 | 0.517 | -58.9% | 0.500 | 0.091 | 0.521 | -59.2% |
| **Inc. R-A** | no | 100 | XL | S | CNN | 0.345 | 0.350 | 0.011 | 0.585 | -41.0% | 0.575 | 0.038 | 0.452 | -23.7% |
| **L-H** | partial | 1000 | XL | S | CNN | 0.180 | 0.176 | 0.017 | 0.231 | -22.0% | 0.232 | 0.008 | 0.408 | -55.9% |

Table S7: Overview of the best and worst TL settings according to MAE for each model type and experiment. We indicate the datasets (Experiment), the DL models, the TL optimality setting, the number of time series (TS), the settings for IC and KP sampling, the MAE results for the simulation-based TL (TL MAE) and baseline DL (DL MAE), the relative change between DL baseline and simulation-based TL (DL Rel.Ch.), the MAE for the ODE calibration baseline (ODE MAE) and the relative change between ODE calibration baseline and simulation-based TL (ODE Rel.Ch.).

| Experiment | Models | TL setting | TS | IC | KP | TL MAE | MAE_std | DL MAE | DL MAE_std | DL Rel.Ch. | ODE MAE | ODE Rel.Ch. |
|---|---|---|---|---|---|---|---|---|---|---|---|---|
| COV-19 | CNN | best | 1000 | XL | L | 1316.006 | 382.725 | 21902.208 | 3741.171 | -0.940 | 1.08e+05 | -0.988 |
| COV-19 | CNN | worst | 1000 | XL | S | 10527.544 | 2997.156 | 21902.208 | 3741.171 | -0.519 | 1.08e+05 | -0.903 |
| COV-19 | DNN | best | 1000 | XL | L | 1188.968 | 271.513 | 11123.954 | 3596.460 | -0.893 | 1.08e+05 | -0.989 |
| COV-19 | DNN | worst | 100 | XL | S | 14550.923 | 4203.451 | 11123.954 | 3596.460 | 0.308 | 1.08e+05 | -0.866 |
| COV-19 | GRU | best | 1000 | L | M | 2243.353 | 1101.747 | 15021.243 | 2086.599 | -0.851 | 1.08e+05 | -0.979 |
| COV-19 | GRU | worst | 10 | L | S | 9591.951 | 2684.640 | 15021.243 | 2086.599 | -0.361 | 1.08e+05 | -0.912 |
| COV-19 | LSTM | best | 1000 | XL | XL | 1532.887 | 559.440 | 19379.778 | 1400.162 | -0.921 | 1.08e+05 | -0.986 |
| COV-19 | LSTM | worst | 10 | M | S | 11837.127 | 811.733 | 19379.778 | 1400.162 | -0.389 | 1.08e+05 | -0.891 |
| Coh. R-A | CNN | best | 10 | S | XL | 10.732 | 0.808 | 12.563 | 0.617 | -0.146 | 10.213 | 0.051 |
| Coh. R-A | CNN | worst | 1000 | S | S | 17.896 | 0.704 | 12.563 | 0.617 | 0.424 | 10.213 | 0.752 |
| Coh. R-A | DNN | best | 10 | XL | L | 10.862 | 1.451 | 12.730 | 0.736 | -0.147 | 10.213 | 0.064 |
| Coh. R-A | DNN | worst | 1000 | XL | S | 17.360 | 0.722 | 12.730 | 0.736 | 0.364 | 10.213 | 0.700 |
| Coh. R-A | GRU | best | 100 | L | L | 9.848 | 0.452 | 12.833 | 0.295 | -0.233 | 10.213 | -0.036 |
| Coh. R-A | GRU | worst | 100 | M | S | 17.114 | 0.418 | 12.833 | 0.295 | 0.334 | 10.213 | 0.676 |
| Coh. R-A | LSTM | best | 100 | M | XL | 9.828 | 0.394 | 12.838 | 0.246 | -0.234 | 10.213 | -0.038 |
| Coh. R-A | LSTM | worst | 100 | S | S | 16.884 | 0.512 | 12.838 | 0.246 | 0.315 | 10.213 | 0.653 |
| Incoh. R-A | CNN | best | 1000 | S | S | 13.339 | 0.348 | 11.972 | 0.207 | 0.114 | 32.223 | -0.586 |
| Incoh. R-A | CNN | worst | 1000 | S | XL | 23.606 | 0.627 | 11.972 | 0.207 | 0.972 | 32.223 | -0.267 |
| Incoh. R-A | DNN | best | 1000 | S | S | 13.002 | 0.288 | 12.042 | 0.167 | 0.080 | 32.223 | -0.597 |
| Incoh. R-A | DNN | worst | 1000 | L | XL | 20.539 | 0.812 | 12.042 | 0.167 | 0.706 | 32.223 | -0.363 |
| Incoh. R-A | GRU | best | 1000 | M | S | 13.306 | 0.154 | 12.320 | 0.110 | 0.080 | 32.223 | -0.587 |
| Incoh. R-A | GRU | worst | 1000 | XL | XL | 20.418 | 0.906 | 12.320 | 0.110 | 0.657 | 32.223 | -0.366 |
| Incoh. R-A | LSTM | best | 1 | S | S | 13.030 | 0.119 | 12.336 | 0.119 | 0.056 | 32.223 | -0.596 |
| Incoh. R-A | LSTM | worst | 1000 | XL | XL | 20.543 | 0.495 | 12.336 | 0.119 | 0.665 | 32.223 | -0.362 |
| L-H | CNN | best | 1000 | M | S | 14.870 | 0.966 | 18.751 | 0.851 | -0.207 | 15.288 | -0.027 |
| L-H | CNN | worst | 1000 | L | XL | 21.970 | 2.215 | 18.751 | 0.851 | 0.172 | 15.288 | 0.437 |
| L-H | DNN | best | 1 | L | XL | 14.878 | 0.990 | 18.509 | 1.038 | -0.196 | 15.288 | -0.027 |
| L-H | DNN | worst | 1000 | M | XL | 25.447 | 1.956 | 18.509 | 1.038 | 0.375 | 15.288 | 0.665 |
| L-H | GRU | best | 1 | XL | L | 15.518 | 1.374 | 18.382 | 0.506 | -0.156 | 15.288 | 0.015 |
| L-H | GRU | worst | 1000 | XL | XL | 21.668 | 1.465 | 18.382 | 0.506 | 0.179 | 15.288 | 0.417 |
| L-H | LSTM | best | 1 | L | XL | 15.218 | 0.910 | 18.328 | 0.678 | -0.170 | 15.288 | -0.005 |
| L-H | LSTM | worst | 1000 | L | XL | 26.084 | 2.300 | 18.328 | 0.678 | 0.423 | 15.288 | 0.706 |

Table S8: Overview of the best and worst TL settings according to 1-PTA for each model type and experiment. *Due to space, we indicate 1-PTA values with PTA in the column headers. We indicate the datasets (Experiment), the DL models, the TL optimality setting, the number of time series (TS), the settings for IC and KP sampling, the 1-PTA results for the simulation-based TL (TL 1-PTA) and baseline DL (DL 1-PTA), the relative change between DL baseline and simulation-based TL (DL Rel.Ch.), the 1-PTA for the ODE calibration baseline (ODE 1-PTA) and the relative change between ODE calibration baseline and simulation-based TL (ODE Rel.Ch.).

| Experiment | Models | TL setting | TS | IC | KP | TL 1-PTA | 1-PTA_std | DL 1-PTA | DL 1-PTA_std | DL Rel.Ch. | ODE 1-PTA | ODE Rel.Ch. |
|---|---|---|---|---|---|---|---|---|---|---|---|---|
| COV-19 | CNN | best | 1000 | M | L | 0.077 | 0.068 | 0.564 | 0.077 | -0.864 | 0.981 | -0.922 |
| COV-19 | CNN | worst | 100 | XL | S | 0.659 | 0.228 | 0.564 | 0.077 | 0.168 | 0.981 | -0.328 |
| COV-19 | DNN | best | 100 | XL | M | 0.035 | 0.031 | 0.478 | 0.064 | -0.928 | 0.981 | -0.965 |
| COV-19 | DNN | worst | 100 | S | S | 0.599 | 0.141 | 0.478 | 0.064 | 0.252 | 0.981 | -0.390 |
| COV-19 | GRU | best | 100 | S | XL | 0.019 | 0.000 | 0.599 | 0.138 | -0.968 | 0.981 | -0.980 |
| COV-19 | GRU | worst | 1 | M | S | 0.347 | 0.194 | 0.599 | 0.138 | -0.420 | 0.981 | -0.646 |
| COV-19 | LSTM | best | 10 | S | L | 0.018 | 0.003 | 0.538 | 0.110 | -0.967 | 0.981 | -0.982 |
| COV-19 | LSTM | worst | 1 | S | M | 0.332 | 0.148 | 0.538 | 0.110 | -0.383 | 0.981 | -0.661 |
| Coh. R-A | CNN | best | 1 | XL | S | 0.237 | 0.032 | 0.496 | 0.060 | -0.521 | 0.521 | -0.544 |
| Coh. R-A | CNN | worst | 100 | L | S | 0.383 | 0.024 | 0.496 | 0.060 | -0.227 | 0.521 | -0.264 |
| Coh. R-A | DNN | best | 1 | M | S | 0.221 | 0.024 | 0.517 | 0.045 | -0.573 | 0.521 | -0.576 |
| Coh. R-A | DNN | worst | 100 | L | S | 0.362 | 0.052 | 0.517 | 0.045 | -0.298 | 0.521 | -0.304 |
| Coh. R-A | GRU | best | 10 | XL | L | 0.212 | 0.017 | 0.517 | 0.091 | -0.589 | 0.521 | -0.592 |
| Coh. R-A | GRU | worst | 100 | M | S | 0.321 | 0.024 | 0.517 | 0.091 | -0.379 | 0.521 | -0.384 |
| Coh. R-A | LSTM | best | 10 | L | M | 0.229 | 0.033 | 0.492 | 0.062 | -0.534 | 0.521 | -0.560 |
| Coh. R-A | LSTM | worst | 1000 | L | L | 0.312 | 0.015 | 0.492 | 0.062 | -0.364 | 0.521 | -0.400 |
| Incoh. R-A | CNN | best | 100 | XL | M | 0.345 | 0.011 | 0.585 | 0.038 | -0.410 | 0.452 | -0.237 |
| Incoh. R-A | CNN | worst | 1 | S | XL | 0.420 | 0.041 | 0.585 | 0.038 | -0.282 | 0.452 | -0.072 |
| Incoh. R-A | DNN | best | 10 | L | M | 0.360 | 0.014 | 0.565 | 0.058 | -0.363 | 0.452 | -0.204 |
| Incoh. R-A | DNN | worst | 100 | S | M | 0.431 | 0.012 | 0.565 | 0.058 | -0.237 | 0.452 | -0.047 |
| Incoh. R-A | GRU | best | 1 | L | L | 0.375 | 0.018 | 0.560 | 0.034 | -0.330 | 0.452 | -0.171 |
| Incoh. R-A | GRU | worst | 10 | M | XL | 0.485 | 0.052 | 0.560 | 0.034 | -0.134 | 0.452 | 0.072 |
| Incoh. R-A | LSTM | best | 1000 | XL | L | 0.360 | 0.049 | 0.555 | 0.041 | -0.351 | 0.452 | -0.204 |
| Incoh. R-A | LSTM | worst | 100 | S | M | 0.475 | 0.054 | 0.555 | 0.041 | -0.144 | 0.452 | 0.050 |
| L-H | CNN | best | 1000 | XL | S | 0.180 | 0.017 | 0.231 | 0.008 | -0.220 | 0.408 | -0.559 |
| L-H | CNN | worst | 1000 | S | M | 0.354 | 0.030 | 0.231 | 0.008 | 0.535 | 0.408 | -0.132 |
| L-H | DNN | best | 100 | M | S | 0.190 | 0.022 | 0.229 | 0.032 | -0.170 | 0.408 | -0.535 |
| L-H | DNN | worst | 1000 | S | XL | 0.448 | 0.016 | 0.229 | 0.032 | 0.958 | 0.408 | 0.097 |
| L-H | GRU | best | 1 | S | XL | 0.310 | 0.071 | 0.246 | 0.018 | 0.260 | 0.408 | -0.241 |
| L-H | GRU | worst | 10 | XL | M | 0.451 | 0.096 | 0.246 | 0.018 | 0.835 | 0.408 | 0.105 |
| L-H | LSTM | best | 1 | S | XL | 0.318 | 0.036 | 0.240 | 0.016 | 0.328 | 0.408 | -0.220 |
| L-H | LSTM | worst | 1000 | L | XL | 0.499 | 0.035 | 0.240 | 0.016 | 1.080 | 0.408 | 0.222 |

### A.9 Visualization of the Time Series Predictions

Besides a quantitative comparison of the time series predictions in the simulation-based TL, DL baseline and ODE baseline settings, we here provide a visualization of their predictions. Figures S7, S8, S9 and S10 (which is identical to Fig. fig:lynx-hare-predictions from the main text) show the predictions of the best performing simulation-based TL model, the DL baseline and ODE baseline on the COVID-19, coherent rotifers-algae, incoherent rotifers-algae and lynx and hares dataset according to Table S4. Predictions are not averaged over seeds.

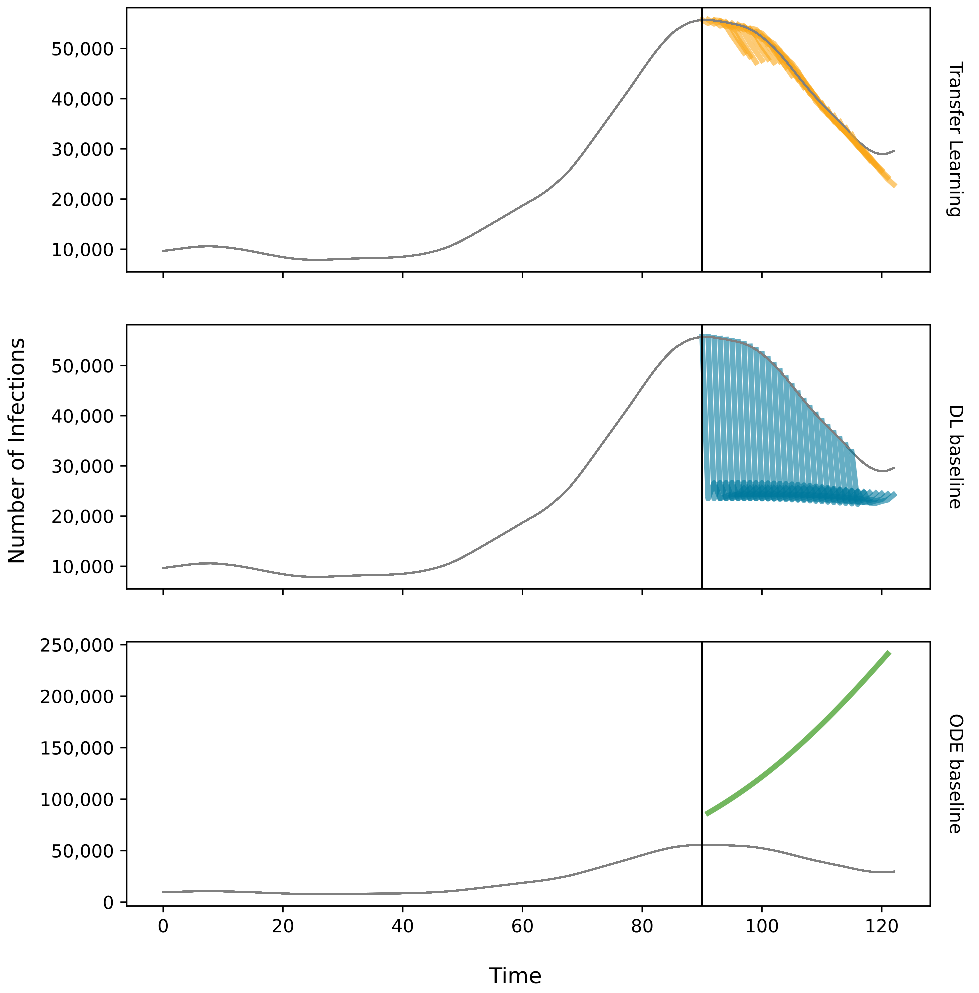

Figure S7: Predicted number of infections on the COVID-19 dataset for the best simulation-based TL setting, DL baseline and ODE baseline. The vertical lines indicate the train-test cutoff for training the TL and DL models. 10% of the train set were used as a validation set. The ODE baseline was fitted to all data points in the train set. It can be observed that only the TL model manages to correctly infer the decline in infections numbers. Fitting the SIR model to the train set leads to an extrapolation of the exponential increase of infections. In conlusion, this explains the findings from Table S4 that the simulation-based TL setting outperforms both baselines by over 90% on MAE.

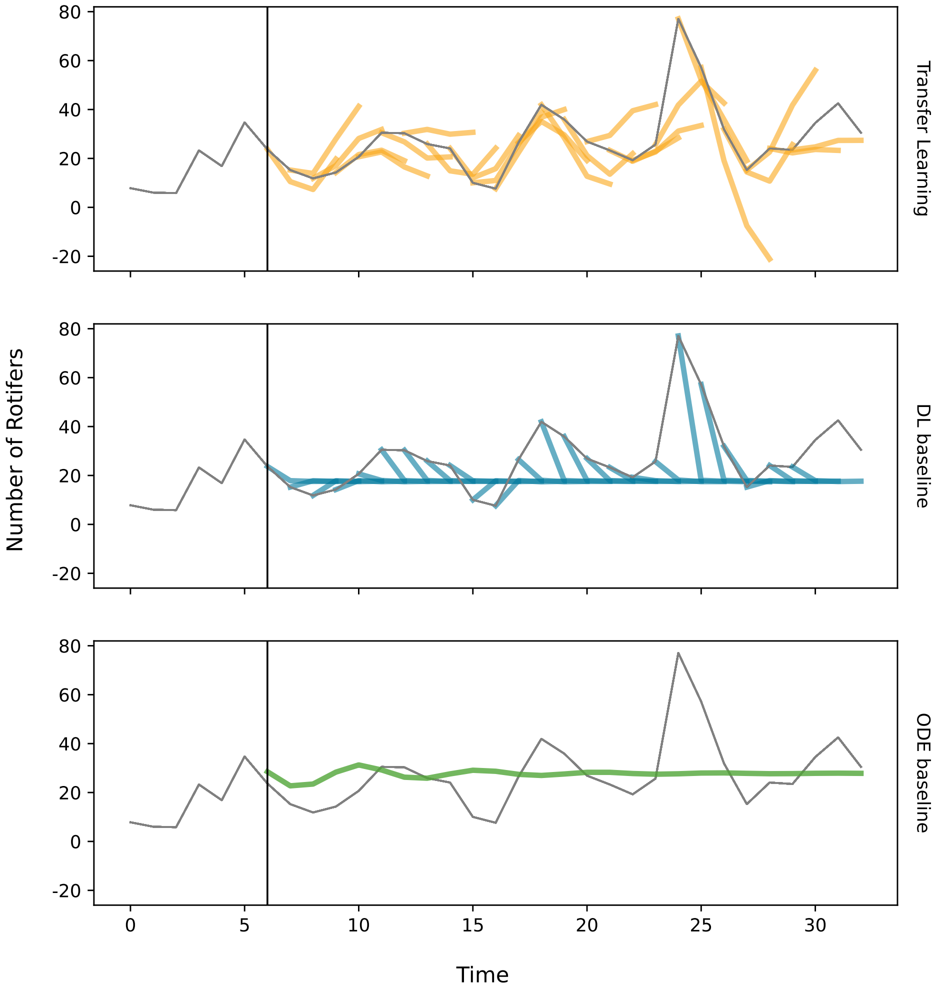

Figure S8: Predicted number of rotifers of the coherent rotifers-algae datasets for the best simulation-based TL setting, DL baseline and ODE baseline. The vertical lines indicate the train-test cutoff used when training the TL and DL models. 10% of the train set were used as a validation set. The ODE baseline was fitted to all data points in the train set. It can be observed that pre-training on synthetic rotifers-algae popluation time series allows the TL model to successfully reproduce the periodic behavior characteristic of the rotifer population development. Despite achieving an adequate fit to the training data for the rotifer and algae populations (see Fig. A.7), the calibrated ODE model fails to accurately capture the subsequent dynamics. Similarly, the limited quantity of training data causes DL baseline model to predict values close to the mean of the training set only.

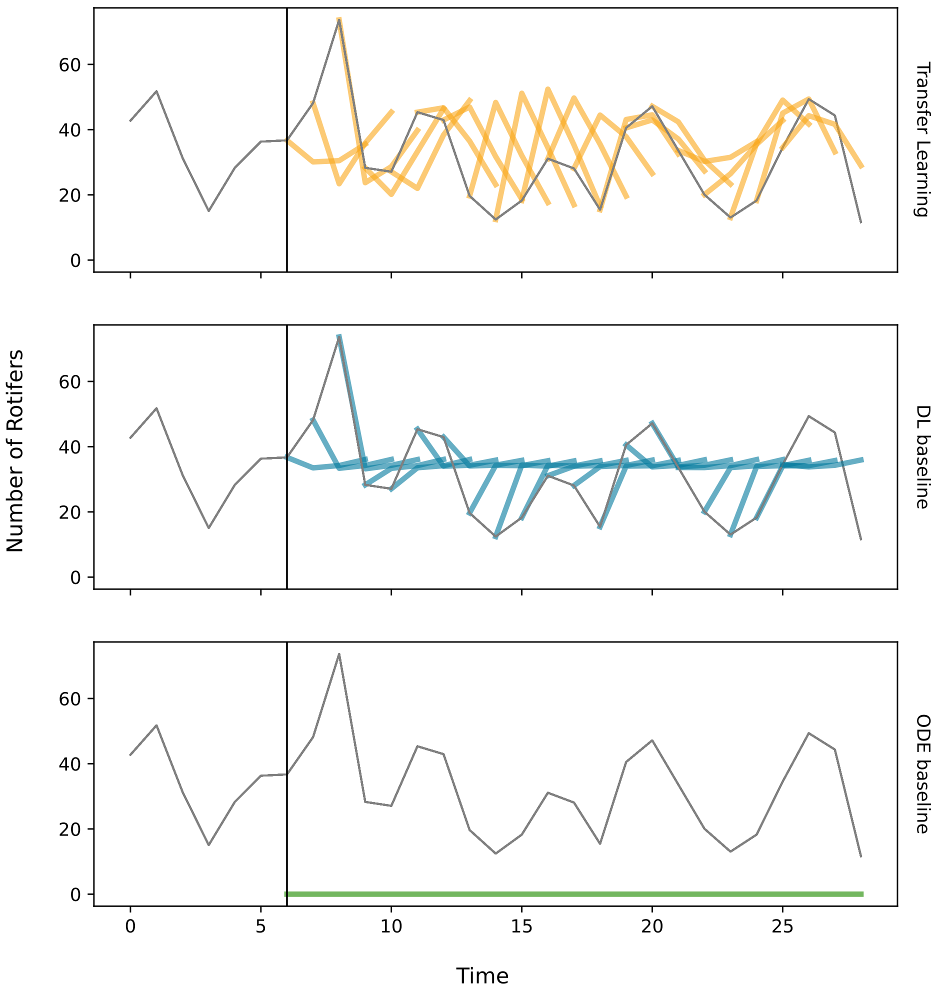

Figure S9: Predicted number of rotifers of the coherent rotifers-algae datasets for the best simulation-based TL setting, DL baseline and ODE baseline. The vertical lines indicate the train-test cutoff used when training the TL and DL models. 10% of the train set were used as a validation set. The ODE baseline was fitted to all data points in the train set. In contrast to Figure S8, none of the three approaches isable to correctly infer the rotifer's population dynamics in the test set. This can be attributed to the inconsistent relation between the rotifer and algae population in this dataset as described in section 2.3. This also provides reasoning for the insufficient fit of the ODE model to the training data.

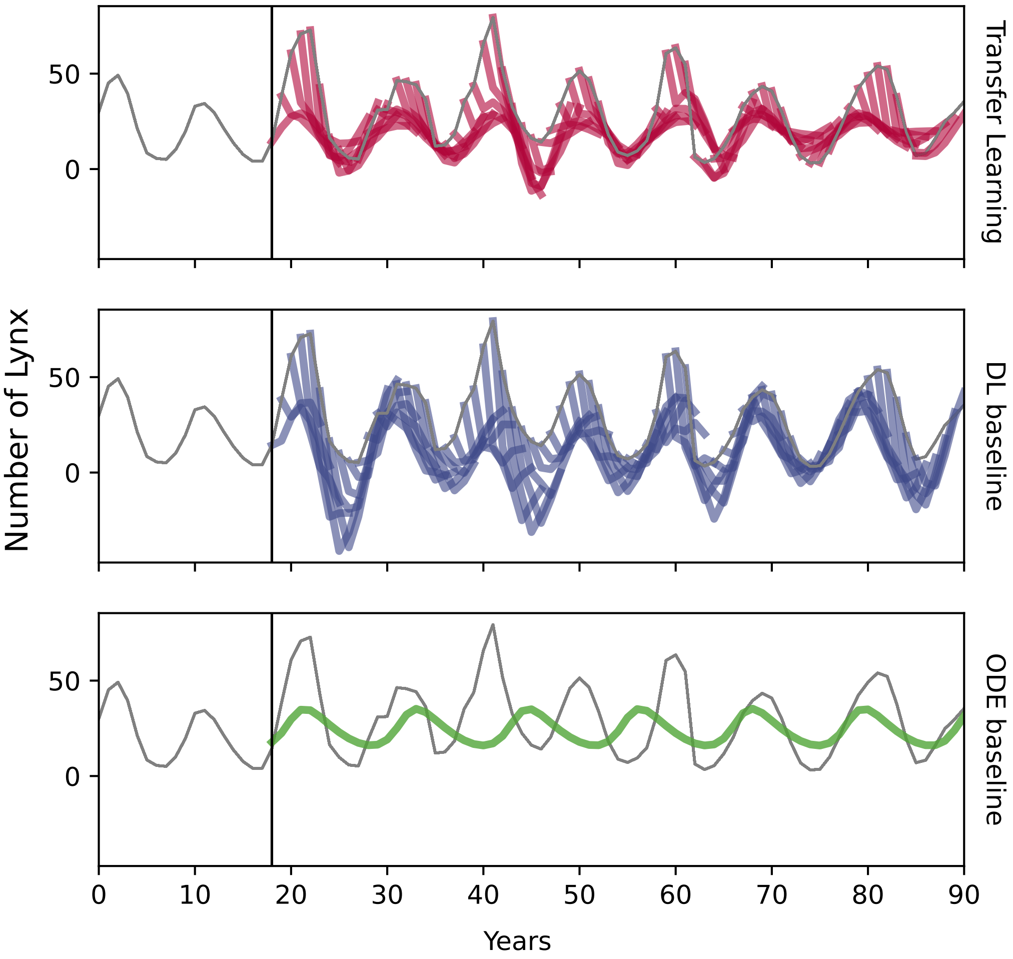

Figure S10: Predicted number of lynx for the best simulation-based TL setting, DL baseline and ODE baseline. This figure is identical to Fig. 2 from the main text and is shown here for sake of completeness. The vertical lines indicate the train-test cutoff used when training the TL and DL models. 10% of the train set were used as a validation set. The ODE baseline was fitted to all data points in the train set. It can be observed that both the TL and DL model pick up the oscillatory trend of the lynx population. However, pre-training on generated time series from the ODE model allows the ML model to substantially reduce the offset between predictions and ground truth. This is in line with the comparison of the quantitative performances in Table S4.

### A.10 Multivariate Analysis of the Impact of Dataset Size and Diversity

### A.11 Impact of Synthetic Dataset Size

Regarding 1-PTA results with the small network architecture, we observe the greatest improvements in the COVID-19 experiments when increasing the synthetic dataset size from 1 to 10 (see Figure S12). Furthermore, we observe that pre-training on any synthetic dataset configuration with 10, 100, or 1000 time series yields near-perfect forecasts regarding 1-PTA. In contrast to the COVID-19 experiment, increasing the synthetic dataset size in the coherent and incoherent rotifers-algae experiments has a smaller impact on 1-PTA scores. In the lynx and hares experiment, we notice a higher variance in TL results for larger synthetic datasets, a trend that is also evident in MAE scores.

For the larger DL model architecture, we make similar observations as for the smaller architectures. In Figure S13, we observe the overall trend that larger synthetic datasets yield better MAE performance of TL runs if time series dynamics in synthetic and real-world datasets are coherent (see COVID-19 and Rotifers and Algae (coherent)). For the incoherent rotifers-algae and lynx and hares, the opposite effect is observed. Larger synthetic datasets improve TL results regarding 1-PTA scores on the COVID-19 and coherent rotifers-algae dataset.

### A.12 Impact of Synthetic Dataset Diversity

#### A.12.1 Univariate Analysis of the Effect of Kinetic Parameters

In comparison to MAE, the length of KP sampling intervals has a smaller influence on the 1-PTA performance of TL runs (see Figure S14). Deteriorating median 1-PTA scores with longer sampling intervals can only be observed for the lynx and hares experiment. Furthermore, we do not observe a clear trend in the variance of 1-PTA scores for longer KP intervals as it is present for MAE scores. The impact of the KP sampling intervals on the variance of MAE and 1-PTA scores is stronger for the large DL model architectures (see Figure S15).

#### A.12.2 Univariate Analysis of the Effect of Initial Conditions

Manipulating dataset diversity by altering the IC sampling intervals shows decisively less influence on simulation-based TL prediction performance, both in terms of MAE and 1-PTA scores, compared to the size of synthetic datasets and KPs. This applies for the small and large DL architectures as shown in Figure S16 and Figure S17, respectively. This observation can be made for the vast majority of DL models across all experiments, as shown in Table S9.

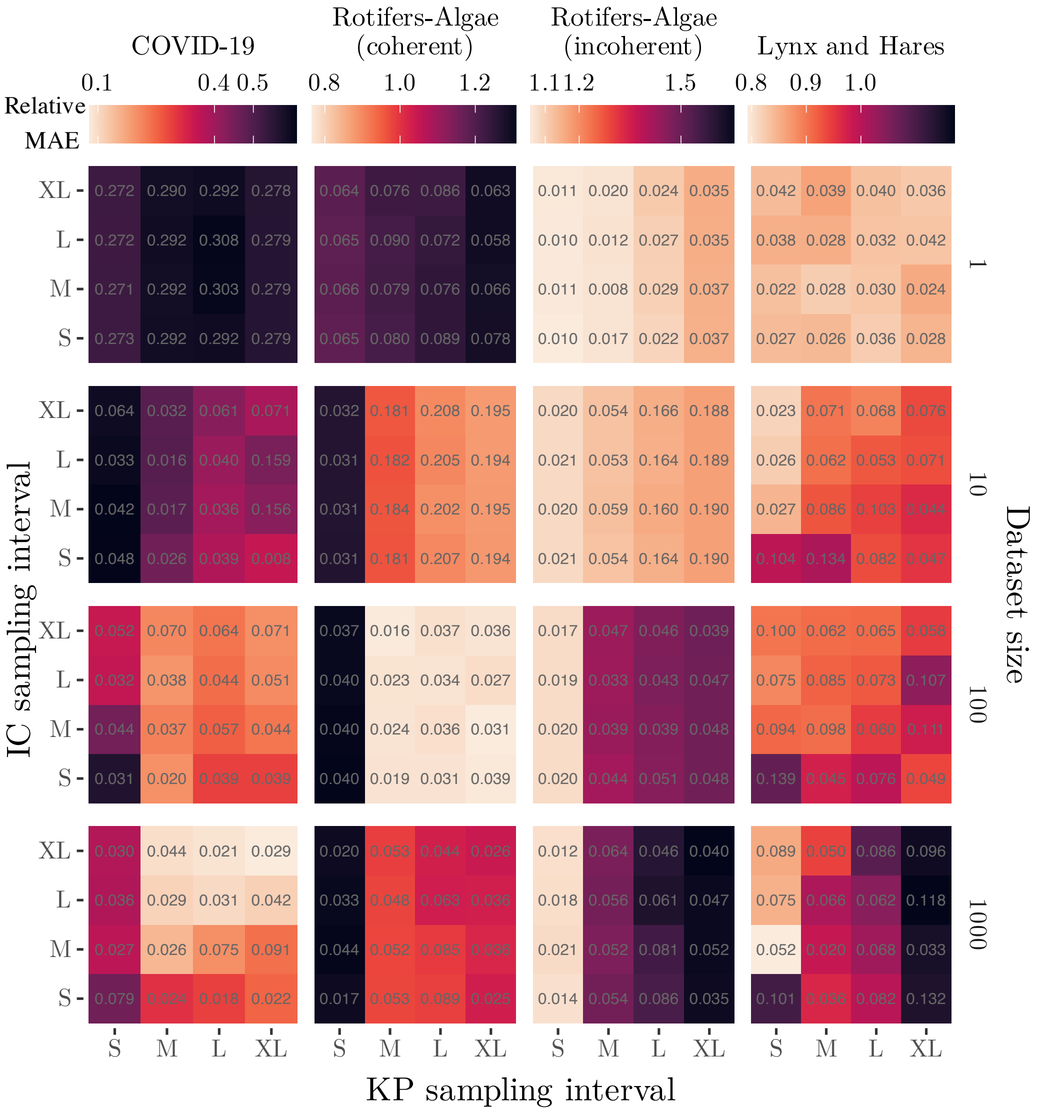

Figure S11: Multivariate analysis of the impact of dataset size, IC-based and KP-based dataset diversity on simulation-based TL performance. For each dataset, we show the MAE for the best-performing DL model type (see Table 2, 'Model' column) relative to its performance in the DL baseline (color-encoded). The standard deviation normalized by the deep learning baseline is shown as number in the corresponding tile. We observe that the optimal choice of the characteristics of synthetic datasets highly varies between real-world target datasets. This underlines the necessity of optimizing ODE-derived synthetic datasets to fully leverage the predictive power of simulation-based TL.

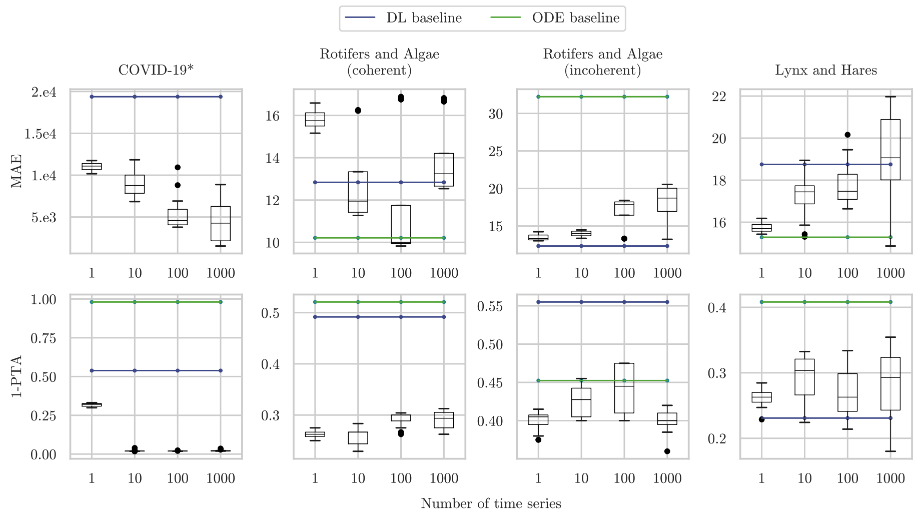

Figure S12: Univariate analysis of how synthetic datasets size impacts TL performance of the *small* DL model architectures. The boxplots visualize the distribution of TL runs ($C \times P$) for each dataset size. TL (black) and DL baseline runs (blue) are shown for the best performing DL model type regarding MAE (see Table 2 in paper). *We exclude the MAE performance ($1.08 \times 10^5$) of the ODE baseline (green) in the COVID-19 experiment to improve scaling. Regarding MAE, larger synthetic datasets improve TL results only if dynamics between the synthetic and the real-world target dataset are coherent (COVID-19 and coherent rotifers-algae). For 1-PTA, the benefit of choosing larger synthetic datasets can only be observed in the COVID-19 experiment.

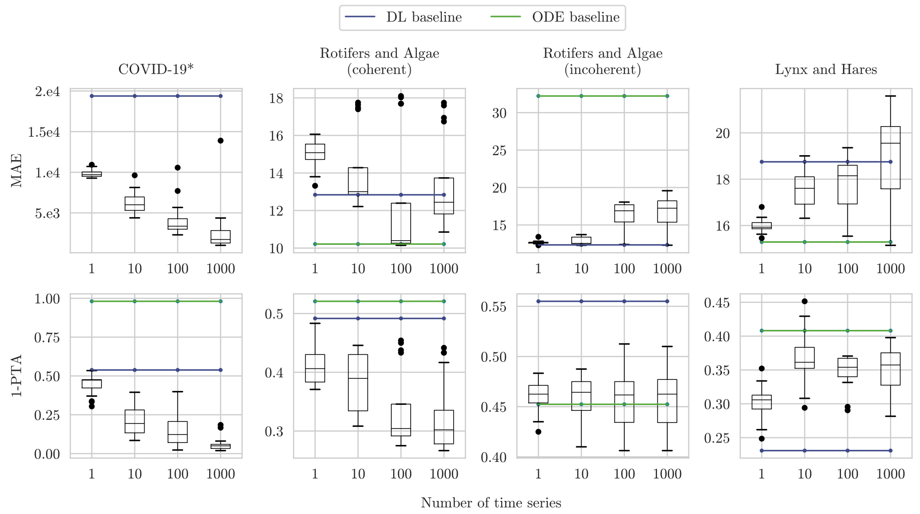

Figure S13: Univariate analysis of how synthetic datasets size impacts TL performance of the *large* DL model architectures. The boxplots visualize the distribution of TL runs ($C \times P$) for each dataset size. TL (black) and DL baseline runs (blue) are shown for the best performing DL model type regarding MAE (see Table 2 in paper). *We exclude the MAE performance ($1.08 \times 10^5$) of the ODE baseline (green) in the COVID-19 experiment to improve scaling. Larger synthetic datasets have a similar effect on TL performance for the large DL architectures as for the smaller ones. However, we observe a greater benefit regarding 1-PTA in the COVID-19 and coherent rotifers-algae datasets when increasing the synthetic dataset size.

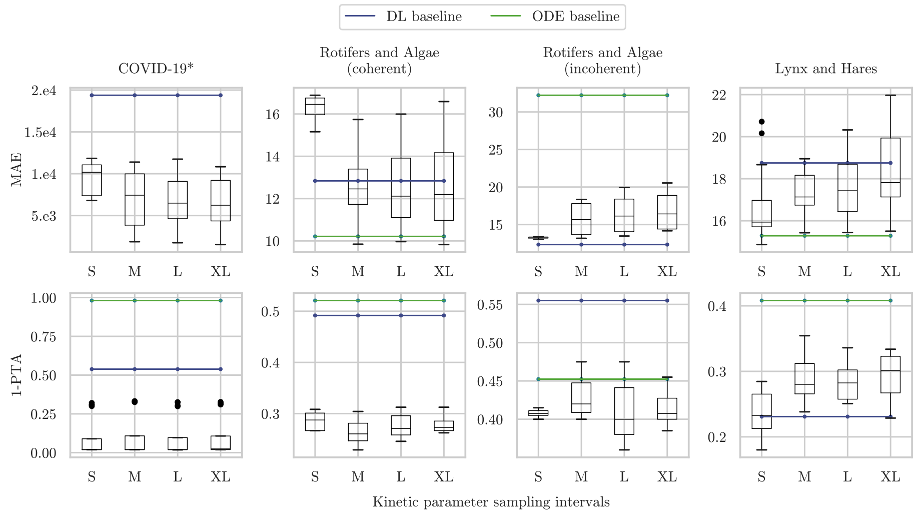

Figure S14: Univariate analysis of how the different KP sampling intervals impacts TL performance of the small DL architectures. The boxplots show the distribution of TL runs ($Z \times C$) for each KP sampling interval size. TL (black) and DL baseline runs (blue) are shown for the best performing DL model (small architecture) regarding MAE (see Table 2). *We exclude the MAE performance ($1.08 \times 10^5$) of the ODE baseline (green) in the COVID-19 experiment to improve scaling. If system dynamics between synthetic and real-world data are coherent, choosing a KP sampling interval greater than S can improve TL results regarding MAE (see COVID-19 and coherent rotifers-algae). The opposite effect is observed if dynamics are incoherent. The 1-PTA scores of TL runs in all experiments are less affected by different-sized KP sampling intervals.

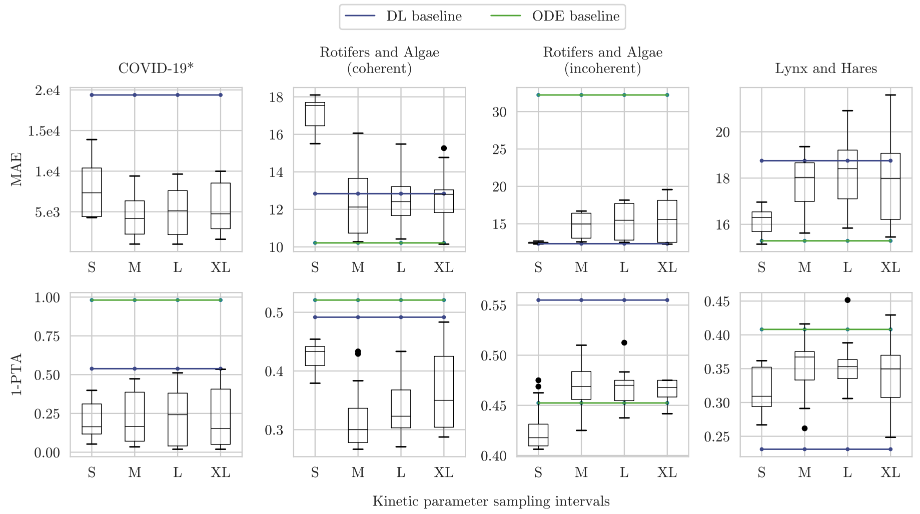

Figure S15: Univariate analysis of how the different KP sampling intervals impact the TL performance of the large DL model architectures. The boxplots visualize the distribution of TL runs ($Z \times C$) for KP sampling interval. TL (black) and DL baseline runs (blue) are shown for the best performing DL model (small architecture) regarding MAE (see Table 2). *We exclude the MAE performance ($1.08 \times 10^5$) of the ODE baseline (green) in the COVID-19 experiment to improve scaling. As for the smaller DL model architectures, increasing the KP sampling intervals has an opposing effect on MAE performance when system dynamics between synthetic and real-world datasets are coherent or incoherent. For the rotifers-algae case studies, this observation can also be made regarding 1-PTA.

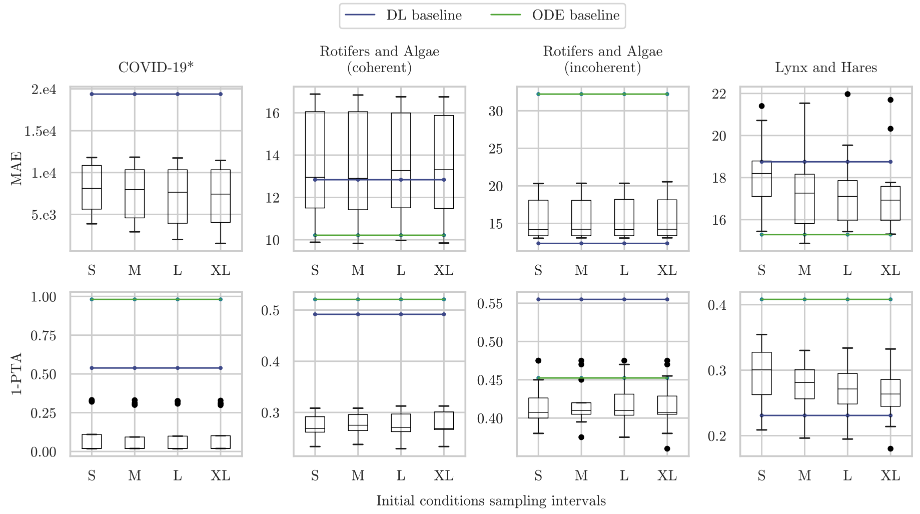

Figure S16: Univariate analysis of how the different IC sampling intervals impact the TL performance for the large DL model architectures. The boxplots visualize the distribution of TL runs ($Z \times P$) for IC sampling interval. TL (black) and DL baseline runs (blue) are shown for the best performing DL model (small architecture) regarding MAE (see Table 2). *We exclude the MAE performance ($1.08 \times 10^5$) of the ODE baseline (green) in the COVID-19 experiment to improve scaling. We observe that different IC sampling interval sizes have a smaller influence on TL performance than synthetic datasets size or KP sampling interval. The shown results of the CNN in the lynx and hares case study are an exception to this finding. This is also indicated in Table S9

.

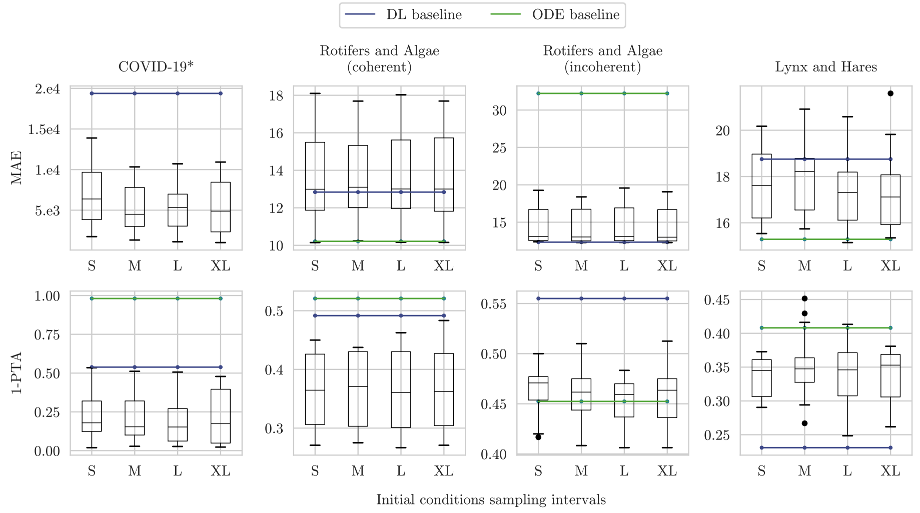

Figure S17: Univariate analysis of how the different IC sampling intervals impact the TL performance for the large DL model architectures. The boxplots visualize the distribution of TL runs ($Z \times P$) for IC sampling interval. TL (black) and DL baseline runs (blue) are shown for the best performing DL model (small architecture) regarding MAE (see Table 2). *We exclude the MAE performance ($1.08 \times 10^5$) of the ODE baseline (green) in the COVID-19 experiment to improve scaling. As for the smaller DL architectures, different IC sampling intervals have a smaller influence on TL results than the choice of datasets size or the KP sampling interval.

### A.13 Impact of Synthetic Dataset Characteristics on Different Deep Learning Model Types

We assess the consistency regarding the impact of different synthetic datasets design choices across DL model types, we qualitatively evaluate the overall observed trend in MAE/1-PTA scores when increasing the dataset size and diversity. We find that different design choices have a similar impact on TL performance across different model types. Please see Table S9 for details. Note that we only consider the small DL architecture for this evaluation.

Table S9: Qualitative analysis on the impact dataset size (TS) and size of IC and KP sampling intervals on the median TL performance. This is shown for each model type in each of the four experiments. ↑ indicates a general increase in median metric scores with more time series or larger IC/KP sampling intervals, ↓ refers to the opposite trend. → represents cases where the median metric scores stay roughly level. We find that the presented findings on the impact of datasets size (A.11), KP (A.12.1) and IC (A.12.2) sampling intervals are overall consistent across the DL model types: For all DL model types, we observe an opposite impact of increasing the synthetic dataset size/KP sampling intervals on MAE and 1-PTA if system dynamics between synthetic and real-world datasets coherent and incoherent.

| | **Exp.** | **L&H** partial inc. | | **Inc. R&A** incoherence | | **Coh. R&A** coherence | | **COV-19** coherence | |
|---|---|---|---|---|---|---|---|---|---|
| | metric | MAE | 1-PTA | MAE | 1-PTA | MAE | 1-PTA | MAE | 1-PTA |
| TS | LSTM | ↑ | ↑ | ↑ | ↑ | ↓ | → | ↓ | ↓ |
| | GRU | ↑ | ↑ | ↑ | → | ↓ | → | ↓ | ↓ |
| | CNN | ↑ | ↑ | ↑ | → | ↓ | ↑ | ↓ | ↓ |
| | DNN | ↑ | ↑ | ↑ | → | ↓ | ↑ | ↓ | ↑ |
| IC | LSTM | → | → | → | → | → | → | → | → |
| | GRU | → | ↑ | → | → | → | → | → | → |
| | CNN | ↓ | ↓ | → | → | → | → | → | → |
| | DNN | → | → | → | → | → | → | → | ↓ |
| KP | LSTM | ↑ | ↑ | ↑ | → | ↓ | → | ↓ | → |
| | GRU | ↑ | ↑ | ↑ | → | ↓ | → | ↓ | → |
| | CNN | ↑ | ↑ | ↑ | → | ↓ | ↓ | ↓ | → |
| | DNN | ↑ | ↑ | ↑ | → | ↓ | → | ↓ | ↓ |

### A.14 Impact of Synthetic Noise

We applied synthetic noise to the pre-training data (for the optimal data characteristics as reported in Table 2 of the main text and S4). The variances for the results on the multiplicative noise from Fig. 5 of the main text are given in Fig. S18.

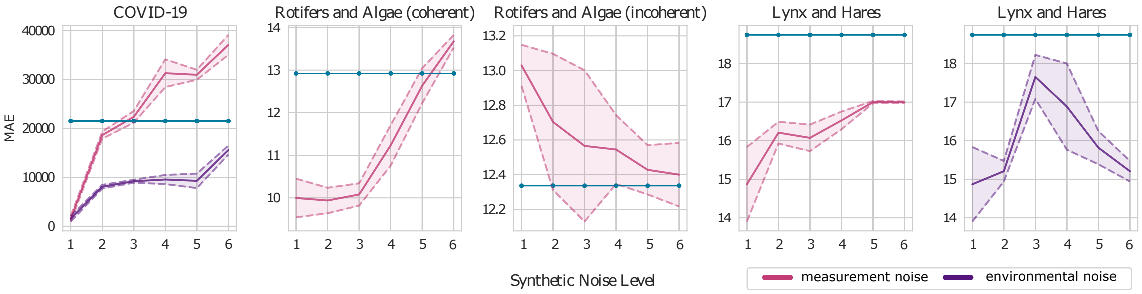

Figure S18: Variances of the univariate analysis of the impact of multiplicative synthetic noise on simulation-based TL performance. Lines show the mean MAE of simulation-based TL performance relative to the DL baseline (blue) for each noise level (see Table S10) over five seeds (as shown in Fig. 5) and their variances. Environmental noise was not used for the rotifers-algae datasets due to numerical instabilities making simulations computationally infeasible.

In addition to multiplicative noise, we also investigate the results when adding additive noise, and show the results for the performance in terms of MAE and 1-PTA (Fig. S19). In Table S10, the different noise levels 1-6 are defined.

We find differences in results for coherent and incoherent settings. Overall, small noise levels seemed beneficial for coherent datasets, whereas larger noise level could provide benefit in incoherent settings.

Table S10: Multiplicative noise levels and dataset-specific noise levels of additive measurement (meas) and environmental (env) noise. The intervals indicate the IQR of the intervals from that noise levels are sampled. The multiplicative noise is sampled from a lognormal distribution with $\mu = 0$ and different standard deviations, the additive noise is sampled from normal distributions with $\mu = 0$ and different standard deviations (see also Table S11). The standard deviations are based on the scale of absolute values in the respective real-world datasets. We do not consider environmental noise when generating synthetic datasets for the rotifers-algae experiments due to numerical instabilities that arose in the SAR ODE system [13] which resulted in infeasible computation times required for simulating sufficiently long time series.

| Noise level | mult. all datasets meas & env | additive COVID-19 meas | env | R-A coh, incoh meas | env | lynx-hares meas | env |
|---|---|---|---|---|---|---|---|
| 1 | 1 | 0 | 0 | 0 | - | 0 | 0 |
| 2 | [0.96,1.04] | [-5.4,5.4] | [-0.674,0.674] | [-2.7,2.7] | - | [-0.084,0.084] | [-0.042,0.042] |
| 3 | [0.92,1.09] | [-43.2,43.2] | [-5.4,5.4] | [-10.8,10.8] | - | [-0.169,0.169] | [-0.084,0.084] |
| 4 | [0.84,1.18] | [-345,345] | [-10.8,10.8] | [-43.2,43.2] | - | [-0.337,0.337] | [-0.169,0.169] |
| 5 | [0.71,1.40] | [-2763,2763] | [-43.2,43.2] | [-172,172] | - | [-0.674,0.674] | [-0.337,0.337] |
| 6 | [0.51,1.96] | [-22102,22102] | [-345,345] | [-691,691] | - | [-1.35,1.35] | [-0.674,0.674] |

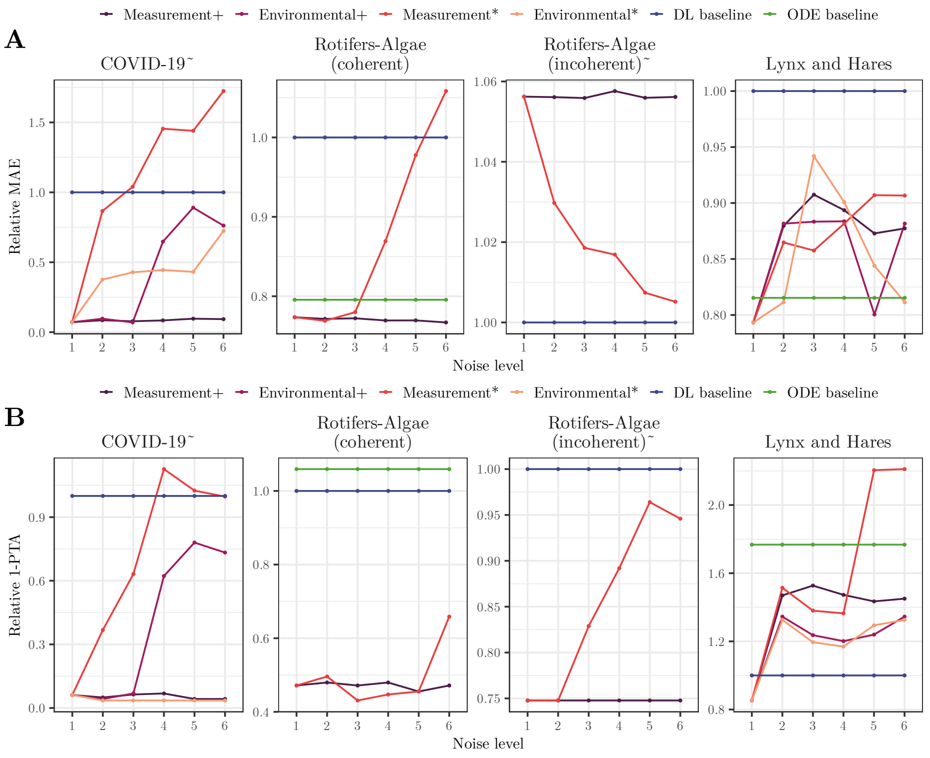

Figure S19: Univariate analysis of the impact of synthetic noise on simulation-based TL performance. Lines show the mean MAE (A) or 1-PTA (B) of simulation-based TL performance relative to the DL baseline (blue) and the ODE baseline (green) for each noise level (see Table S10). Environmental noise was not used for the rotifers-algae datasets due to numerical instabilities making simulations computationally infeasible. ODE baselines are given in green. ˜ We exclude the MAE performance of the ODE baseline in the COVID-19 and incoherent rotifer-algae experiment to improve scaling. Low noise levels are optimal for real-world datasets coherent with synthetic data, while larger noise levels can be beneficial for simulation-based TL in incoherent settings. Additive measurement noise is overall less influential, possibly due to lower effective levels.

### A.15 Synthetic-Only Experiments to Investigate the Relationship Between Noise in Synthetic and Real-World Data and its Impact on Simulation-Based TL Performance

In addition to exploring the benefit of adding noise to synthetic datasets when predicting real-world system dynamics in simulation-based TL, we investigate the hypothesis that similar noise levels in synthetic and real-world datasets improve the prediction performance in a TL setup. To do so, we design synthetic-only experiments using synthetic data generated from the ODE models also for mimicking the real-world, target dataset in which we then can control the noise type and level. Thus, we work in a setting with coherence between pre-training and fine-tuning data (and both datasets are ODE-derived).

For the synthetic-only experiment, next to multiplicative noise that follows a lognormal distribution with noise levels as described in A.1.3, we also consider additive noise. Thereby, we model noise using a normal distribution with $\mu = 0$ and $\sigma$ being defined differently for each ODE model because the impact of additive noise depends on the order of magnitude of data without noise. Details for the standard deviation $\sigma$ used in each experiment are provided in Table S11.

Table S11: ODE-specific standard deviations used to sample six different levels of additive measurement and environmental noise. The standard deviations are based on the scale of absolute values in the respective real-world datasets. We do not consider environmental noise when generating synthetic datasets for the rotifers-algae experiments due to numerical instabilities that arose in the SAR ODE system [13] which resulted in infeasible computation times required for simulating sufficiently long time series.

| ODE Model | Additive Measurement | Additive Environmental |
|---|---|---|
| SIR | 0, 8, 64, 512, 4096, 32768 | 0, 1, 8, 16, 64, 512 |
| SAR | 0, 4, 16, 64, 256, 1024 | - |
| LV | 0, 0.125, 0.25, 0.5, 1, 2 | 0, 0.0625, 0.125, 0.25, 0.5, 1 |

#### A.15.1 Experimental Setup to Investigate the Impact of Noise in Synthetic Datasets on Transfer Learning

We describe the setup to investigate how the noise relationship between pre-training and target dataset influences TL performance. To reduce computational requirements, we consider a setting with intermediate pre-training dataset sizes, i.e., ten time series, and mimick a small data setting by only sampling one target time series (all of length 100). For the synthetic data for pre-training, we generate these ten time series by sampling IC and KP from intervals with $i = 2$ (i.e., L, see Section A.1.2), while we use fixed values ($i = 0$, meaning sampling interval S, see Section A.1.2) for the target time-series. This ensures that pre-training data and target data can differ sufficiently, but they still lie in the same domain, thus guaranteeing the coherence between the dynamics. For noise, we using each combination of noise type (either measurement or environmental), noise operations (either additive or multiplicative), and one of the six noise levels as described above. Please note that to reduce complexity, we only compare simulation-based TL for the same noise types and noise operations between pre-training and target dataset, and only investigate the effect of differences in noise levels.

To yield robust insights, we compute five runs with different seeds for data generation and Machine Learning (ML) training for each experimental setup.

We conduct simulation-based TL experiments for each of the three biological systems (SIR, SAR, LV ODE models) with each combination of two noise types, two noise operations, four DL architectures, and evaluated the performance for two different metrics (MAE, 1-PTA). For each experiment, we investigate $6 \times 6$ noise level settings, i.e., all combinations of the six noise levels for the synthetic and target data. Note that we did not consider environmental noise for the SAR model because it induced instabilities into the system that caused enormously increased runtime for simulating sufficiently long time series.

#### A.15.2 Measuring Similarity Between Noise Levels and Impact on Performance: NPC Measure

To enable direct comparisons for this high-dimensional setting, we aimed to summarize the effect of the similarity between noise levels in the synthetic dataset for pre-training (synthetic noise) and in the target dataset (observed noise) on simulation-based TL performance in a single value for each experiment. This measure should, in principle, retrieve the correlation between similarity in noise levels and simulation-based TL performance. However, assessing the similarity between noise levels is not straightforward, e.g., it is not well-defined whether the difference between noise level 0 and 1 is smaller, larger or equal to the distance between noise level 1 and 2.

This is why we introduce the new Noise-Performance Coherence (NPC) measure that is tailored to our purpose. The NPC measure summarizes the relationship between the similarity in noise during pre-training and noise in fine-tuning target dataset to the simulation-based TL performance. Thereby, a fixed observed noise level in the target data is considered. Thereby, the measure exploits the property that differences of noise levels in one direction are well-ordered (i.e., noise level 2 is less distant from noise level 1 than noise level 3 is from noise level 1). Thus, instead of a single correlation using differences in noise levels, two correlations directly between noise levels and TL performance are computed: (1) those over pre-training datasets with noise levels smaller than or equal to the observed noise, and (2) those over pre-training datasets with noise levels equal to or larger than the observed noise. We expect the first correlation to be negative if TL performance is optimal when synthetic and observed noise are equal (i.e., 1-PTA or MAE are minimal if synthetic noise increases towards observed noise), while the second correlation is expected to be positive in that case (the larger the synthetic noise the further away it is from observed noise, and the worse predictions get). Thus, we combine the two correlations with opposite signs for the final NPC measure (applying a negative sign to the first correlation). Both correlation values are averaged, weighted by the number of datapoints that contribute to each correlation because we consider this number to increase the reliability of the correlation value.

Thus, the NPC measure takes values between -1 and 1. Positive NPC measures indicate that we find better TL performance for similar synthetic and observed noise, negative values indicate the opposite, and values around zero indicate a lack of or non-monotonous relationship.

In mathematical terms, we consider an ordered tuple of $l+1$ noise levels, denoted as $N = (n_0, n_1, \ldots n_l)$ with $n_0 < n_1 < \cdots < n_l$. For computing our NPC measure for an observed noise level $n_k$, we consider a set $P_k$ of pairs of synthetic noise levels and the TL prediction performance when pretraining on synthetic data with that synthetic noise level and transferring to the observed noise level $n_k$. The NPC measure for the observed noise level is defined as the weighted average of the negative Spearman correlation between synthetic noise levels and prediction performances for noise levels smaller or equal to $n_k$, and the Spearman correlation for noise levels greater or equal to $n_k$. Weights are calculated by the number of pairs contributing to each correlation computation, i.e., $k$ as the number of synthetic noise levels smaller than $n_k$ for synthetic noise levels smaller or equal to the observed noise level, and $l-k$ as the number of synthetic noise levels greater than $n_k$ for synthetic noise levels greater or equal to the observed noise level.

The NPC measure for observed noise level $n_k$ is given by:

$$\text{NPC}(n_k) = \frac{(l-k)\rho(S_n^{\uparrow}, S_p^{\uparrow})}{l} - \frac{k\rho(S_n^{\downarrow}, S_p^{\downarrow})}{l}, \tag{13}$$

where $\rho$ is the Spearman correlation and $|N| = l+1$ is the number of elements in the ordered tuple of noise levels $N$, and $S_n^{\uparrow}, S_p^{\uparrow}, S_n^{\downarrow}, S_p^{\downarrow}$ are the projections of the contributing pairs $P_k$ on the synthetic noise levels ($n$) or the TL prediction performances ($p$) stratified by the synthetic noise level being smaller ($\uparrow$) or larger ($\downarrow$) than the observed noise level $n_k$:

$$\begin{aligned} S_n^{\uparrow} &= \{n \mid (n,p) \in P_k \wedge n \geq n_k\} \\ S_p^{\uparrow} &= \{p \mid (n,p) \in P_k \wedge n \geq k\} \\ S_n^{\downarrow} &= \{n \mid (n,p) \in P_k \wedge n \leq k\} \\ S_p^{\downarrow} &= \{p \mid (n,p) \in P_k \wedge n \leq k\} \end{aligned} \tag{14}$$

##### A.15.3 Time Series Forecasting and Training Setup for Synthetic-Only Noise Experiments

The prediction tasks for each biological system, the evaluation and train-test splits are the same as described in Section A.6, except for the coherent rotifers-algae where we use an input and output of five timesteps.

As in the other setups, also for the synthetic-only experiments, the synthetic target data sets are time series of length 100. We restrict fine-tuning to the first 80 time points as train set for the SAR and LV model, and to the first 55 time points as train set for the SIR model. The other time points are used as test set to compute the MAE and 1-PTA (using rolling averages, see Section A.6.1).

In the context of transfer learning, we pre-train our DL models using synthetic data, freeze the feature extraction layer, and fine-tune the fully connected layers for five epochs with the target datasets. A validation set consisting of 10% of synthetic and the real-world train time series is utilized. To prevent overfitting during pre-training, we train all models until the validation loss on the pre-training dataset stabilizes. Throughout the process, we use checkpointing to save the best model state based on validation loss for future tasks.

##### A.15.4 The Impact of the Relation Between Synthetic and Observed Noise

We explore the relationship between noise in the synthetic dataset used for pre-training and the dataset on which the model should be fine-tuned and its effect on simulation-based TL performance.

Overall, as exemplarily shown for the LV predator-prey model with multiplicative noise and the CNN as DL architecture (see Figure S20), we find that there is an influence of the synthetic noise on the prediction performances for most noise conditions in all three ODE models and with all four DL architectures. Exceptions are the experiments using additive environmental noise for the SIR and LV ODEs where, due to a high variance in TL performance across seeds, we cannot see a conclusive influence of the synthetic noise to the prediction performance. Therefore, these noise settings were disregarded in the subsequent analyses.

For the remaining noise settings, we examine the hypothesis whether consistency between the synthetic and observed noise levels is required for optimal simulation-based TL. For that purpose, we use our NPC that is a measure for how strongly the similarity in noise levels and simulation-based TL performances correlate. As shown in Figure S21, NPC tends to have the highest and positive values when the observed dataset does not contain noise (noise level is 0) for the SIR and LV model. This suggests that if the observed, target dataset does not have any noise, we should also use a synthetic dataset without noise. For higher observed noise levels, the NPC is lower but still mostly greater than 0. NPC values $> 0$ suggest that simulation-based TL performance overall improves when synthetic and observed noise level become more similar. Consequently, for higher noise levels, it seems useful to have the same synthetic noise level as the observed noise level. However, the effect is smaller than for low observed noise levels. The same is true for the SIR model when investigating additive measurement noise, where the NPC yields slightly positive values for all observed noise levels. For the SAR model, the NPC measure is mostly slightly positive, indicating that similar noise levels between datasets could improve prediction accuracy. Although negative NPC values appear, which would indicate having synthetic noise levels different from observed noise levels is beneficial for simulation-based TL, negative NPC values are rare and of small magnitude.

Overall, from our experiments, we conclude that having the same noise levels in data for pre-training as well as for fine-tuning can yield better simulation-based TL performance in many cases,

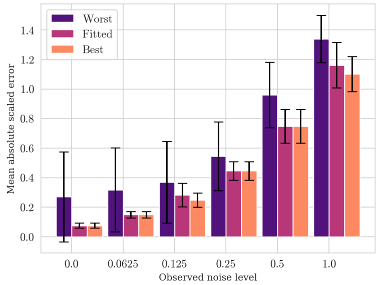

Figure S20: Barplot illustrating the TL performance mean and variance of the CNN over 5 runs of the best-performing (best, orange) and worst-performing (worst, indigo) synthetic noise level per observed noise level for the LV model using multiplicative environmental noise. Performance is indicated in terms of scaled MAE. We also show the performance of the synthetic dataset whose noise level matches the observed noise level (fitted, purple). Differences between the bars for a fixed observed noise level indicate that the synthetic noise has an influence on the simulation-based TL prediction performance.

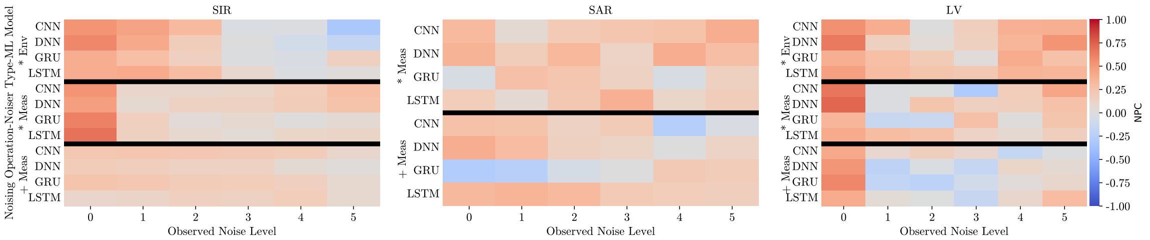

Figure S21: Heatmaps show the NPC as a measure of required consistency between synthetic and observed noise levels in target data for improving TL prediction for each ODE model and each considered noise setting. The black horizontal lines are a visual guidances to separate the noise operations and types. For most cells, especially for observed noise level 0 (no noise), we see an NPC greater than 0 indicating that higher similarity between synthetic and observed noise improves simulation-based TL prediction performance.

but not in all, and it seldom hampers simulation-based TL.

[8] Maximilian Kleissl, Lukas Drews, Benedict B Heyder, Julian Zabbarov, Pascal Iversen, Simon Witzke, Bernhard Y Renard, and Katharina Baum. SimbaML: Connecting mechanistic models and machine learning with augmented data. *ICLR Tiny Paper*, 2023.

[9] A J Lotka. Analytical note on certain rhythmic relations in organic systems. *Proc. Natl. Acad. Sci. U. S. A.*, 6(7):410–415, 1920.

[10] D A MacLulich. *Fluctuations in the numbers of the varying hare (Lepus americanus)*. University of Toronto Press, Toronto, 1937.

[11] Jorge Nocedal and Stephen J Wright. Sequential quadratic programming. In Jorge Nocedal and Stephen J Wright, editors, *Numerical Optimization*, pages 526–573. Springer New York, New York, NY, 1999.

[12] Robert Koch-Institut. SARS-CoV-2 Infektionen in Deutschland, 2022.

[13] Benjamin Rosenbaum, Michael Raatz, Guntram Weithoff, Gregor F Fussmann, and Ursula Gaedke. Estimating parameters from multiple time series of population dynamics using bayesian inference. *Frontiers in Ecology and Evolution*, 6, 2019.

[14] Bao Y Sciscent, Caroline D Eisele, Lisa Ho, Steven D King, Rohit Jain, and Reshma R Golamari. COVID-19 reinfection: the role of natural immunity, vaccines, and variants. *J Community Hosp Intern Med Perspect*, 11(6):733–739, 2021.

[15] Vito Volterra. Fluctuations in the abundance of a species considered mathematically. *Nature*, 118:558–560, 1926.

[16] Cort J Willmott and Kenji Matsuura. Advantages of the mean absolute error (MAE) over the root mean square error (RMSE) in assessing average model performance. *Clim. Res.*, 2005.
